# A microbiota-derived bile acid modulates biofilm formation by the probiotic strain *Escherichia coli* Nissle 1917

**DOI:** 10.1101/2025.06.02.657493

**Authors:** Elena K. Perry, Barath Udayasuryan, Elias K. Zegeye, Christopher M. Rose, Mike Reichelt, Man-Wah Tan

## Abstract

Bacteria that colonize the human gut must withstand a variety of stressors, including detergent-like compounds known as bile acids. Here, we investigated how bile acids found in the human cecum and colon impact the behavior of the probiotic strain *Escherichia coli* Nissle 1917 (EcN). We found that lithocholic acid (LCA), which is a microbiota-derived secondary bile acid, promotes the formation of a distinctive surface-coating biofilm by EcN, including on an organoid-derived model of the human colonic epithelium. Mechanistic investigations revealed that LCA upregulates the production of several components of flagella, which are essential for LCA-induced biofilm formation and form part of the biofilm matrix. Furthermore, LCA-induced biofilm formation helps EcN compete against certain pathogenic strains. Taken together, our findings shed light on how an abundant colonic metabolite influences the behavior of a clinically-proven probiotic strain, triggering the formation of biofilms that may contribute to pathogen suppression.

## INTRODUCTION

Understanding how bacteria behave in their natural context is a fundamental challenge in microbiology. Standard *in vitro* cultures often bear little resemblance to the conditions bacteria face in the wild, whereas *in situ* studies can face issues such as limited accessibility, heterogeneity, and the presence of numerous confounding variables. Nevertheless, controlled experiments have revealed that environmental factors including temperature, flow, and extracellular small molecules can influence bacterial physiology, behavior, and interspecies interactions.^1–3^ For host-associated microbial communities such as the human gut microbiome, uncovering environment-phenotype relationships will likely be crucial to moving from observational studies to the development of microbe-focused strategies to promote health and ameliorate disease. One such strategy that has been under exploration for decades is the use of probiotics, which are defined as live microorganisms that provide health benefits when administered in sufficient quantities.^4^ However, despite the attention that probiotics have received, their practical applications remain limited by inter-individual variability in efficacy and colonization of the gut.^5^ This in turn may reflect our incomplete understanding of how these organisms respond to the complex environment of the human gut.

One of the most-studied probiotic strains is *Escherichia coli* Nissle 1917 (EcN). First isolated by Dr. Alfred Nissle more than 100 years ago from a German soldier who was resistant to infectious diarrhea, EcN has since been used to treat gastrointestinal disorders ranging from acute colitis to chronic constipation and inflammatory bowel diseases.^6^ The molecular characteristics of this unique strain have been extensively investigated in recent decades. Genomic studies have revealed that EcN is closely related to uropathogenic *E. coli* strains belonging to the O6 serotype.^7^ However, while EcN possesses important fitness factors such as iron acquisition systems, it lacks classical pathogenicity factors, such as α-hemolysin.^7^ It is also sensitive to serum due to expressing an unusual semi-rough form of lipopolysaccharide (LPS), in which the O-antigen is truncated after the first residue due to a nonsense mutation in the O-antigen polymerase gene.^8^ Together, this combination of characteristics likely helps EcN colonize the gut while limiting its potential for systemic pathogenicity or other negative impacts on the host.

Numerous mechanisms have been proposed to explain how EcN promotes the resolution of gastrointestinal disorders, ranging from restriction of pathogenic strains via nutritional competition or the secretion of antimicrobial compounds, to immunomodulatory effects, to strengthening of the intestinal epithelial barrier.^6,9–12^ EcN may also form biofilms on the intestinal mucosa.^13,14^ Biofilms have not yet been directly observed for this strain *in vivo*, but indirect evidence for biofilm formation comes from the fact that the expression of adhesins by EcN is necessary for stable colonization of the murine gut.^13^ Besides promoting persistence in the dynamic environment of the gut and direct interactions with epithelial and immune cells, biofilm formation may help EcN suppress pathogens and prevent invasion of the intestinal epithelium, and thus contribute to its probiotic activity.^14,15^ However, little is known about how the gut environment impacts biofilm formation in EcN.

In several other gut bacteria, biofilm formation is regulated by gut-specific signals, with one of the most commonly implicated signals being bile acids.^16–19^ Bile acids are detergent-like compounds produced by the liver and secreted into the small intestine to aid the digestion and absorption of lipids. After transiting into the cecum and colon, the host-produced bile acids, known as primary bile acids, are transformed by the gut microbiota into numerous derivatives, known as secondary bile acids. Individual bile acids have distinct physicochemical properties and can exert differential effects both on host cells and on bacterial physiology,^20–22^ but the latter effects, including the induction of biofilm formation, are not conserved across species.^17,23^ Moreover, while the adaptation of EcN to the murine gut has been investigated,^24^ the environment of the human colon differs from that of the mouse in several respects, including bile acid composition,^25^ leaving the response of EcN to human bile acids unexplored.

In this study, we hypothesized that biofilm formation by EcN would be influenced by the bile acids that predominate in humans, and that different bile acids would have different effects. We found that a specific secondary bile acid, lithocholic acid (LCA), triggers the formation of a distinctive type of biofilm by EcN at physiologically-relevant concentrations, both on abiotic surfaces and on an organoid-derived *in vitro* model of the human intestinal epithelium. We show that LCA-induced biofilm formation is independent of several well-studied *E. coli* biofilm factors, including curli, but that LCA upregulates the expression of flagellar components and flagella appear to play a critical role in the biofilm extracellular matrix. Furthermore, LCA-induced biofilm formation is strain-specific and can increase EcN’s competitive fitness during surface colonization. Taken together, our findings shed light on how an abundant colonic metabolite alters the behavior of a clinically-proven probiotic strain, promoting the formation of biofilms that may contribute to colonization resistance against pathogen invasion.

## RESULTS

### Lithocholic acid triggers formation of a surface-coating biofilm by EcN

To investigate whether bile acids affect EcN biofilm formation under conditions that mimic the human gastrointestinal tract, we grew EcN anaerobically in YCFA medium individually spiked with nine of the major bile acids found in the human gastrointestinal tract (Fig. 1). We chose YCFA as our baseline condition because unlike other media in which EcN biofilm formation has previously been studied (namely, LB broth and minimal media),^13–15^ YCFA supports the growth of a broad range of human gut bacterial species,^26^ suggesting that it better represents the nutritional environment of the human colon. Under this condition, the secondary bile acid lithocholic acid (LCA) triggered a dramatic increase in crystal violet staining starting at 25 µM (Fig. 2A-B). The other tested bile acids showed no effect on EcN biofilm formation up to at least 100 µM. At higher concentrations, starting at 250 µM and 500 µM respectively, chenodeoxycholic acid (CDCA) and ursodeoxycholic acid (UDCA) also stimulated biofilm formation (Supplementary Fig. 1A). However, these concentrations far exceed what has been reported for these bile acids in healthy human colon contents, whereas 25-100 µM spans the range reported for LCA (Supplementary Table 1).^27,28^ None of the bile acids were toxic to EcN up to 500 µM (Supplementary Fig. 1B). In addition, when combined with other non-biofilm-inducing bile acids, LCA still triggered formation of the surface-coating biofilm (Supplementary Fig. 1C).

**Fig. 1:**
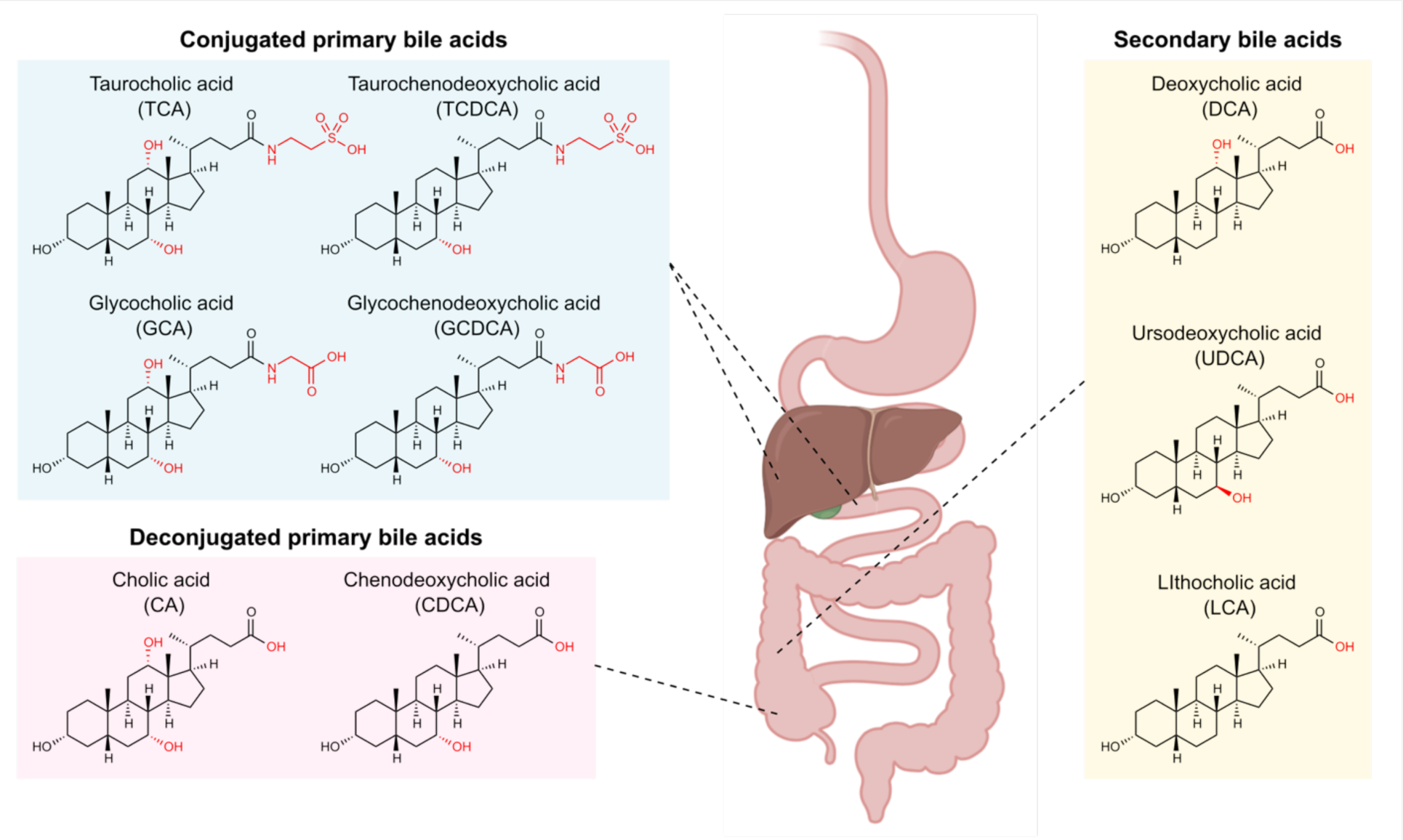
Structures of the nine bile acids tested in this study. The variable functional groups are shown in red. Dotted lines point to the approximate regions of highest concentration along the GI tract for each category of bile acids. The GI tract schematic was created in Biorender.

**Fig. 2:**
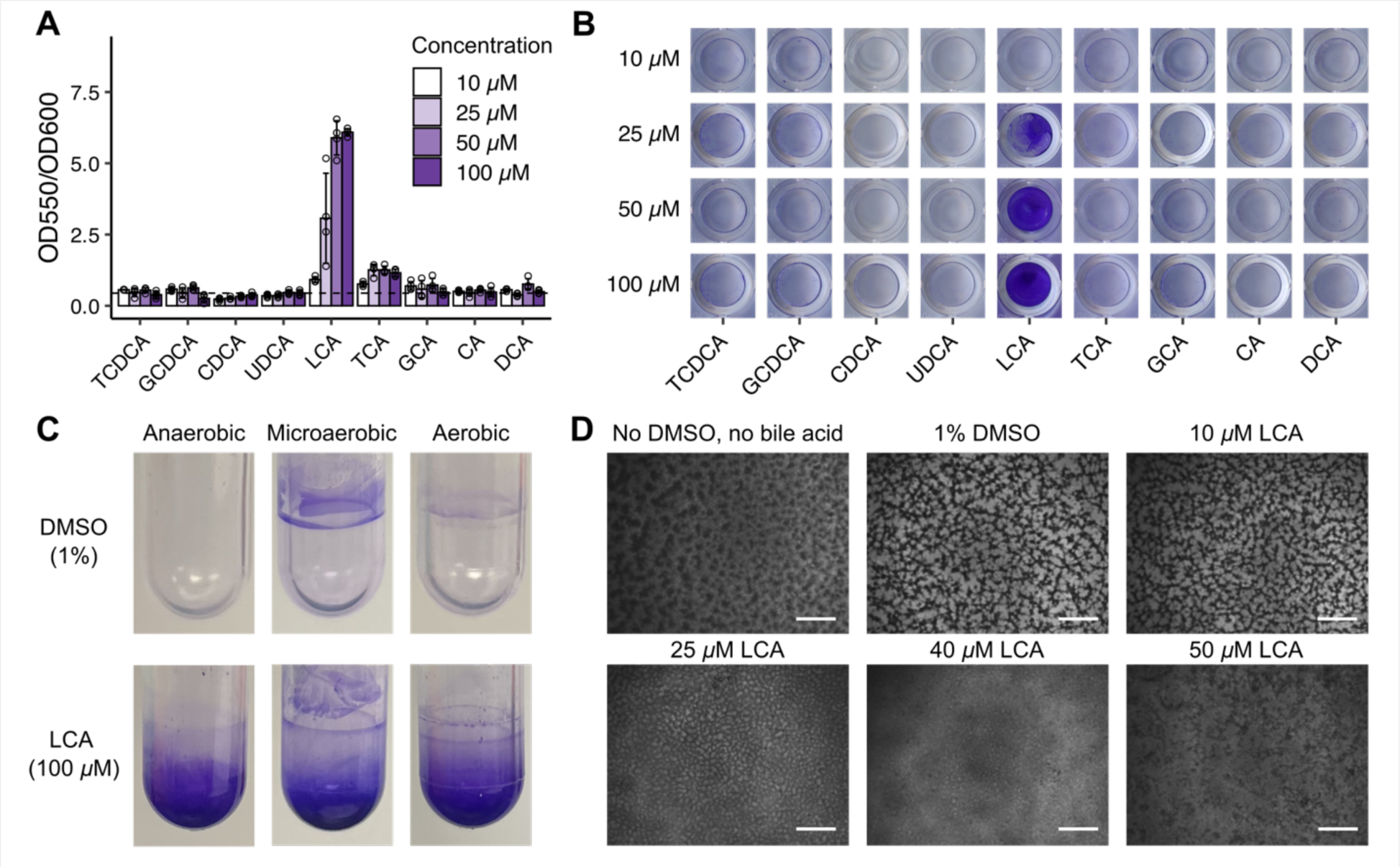
(A) Quantification of biofilm formation (crystal violet absorption at OD_550_ normalized to OD_600_ as a proxy for growth) after 24 hrs of treatment with 10, 25, 50, or 100 µM individual bile acids. Error bars mark the standard deviation (*n* = 4 replicate wells from a representative experiment). (B) Representative images of the wells plotted in (A). (C) Photos of biofilms formed by EcN in 5 mL tube under different oxygen concentrations, in the presence of 1% DMSO (solvent control) or 100 µM LCA. (D) Phase contrast images of aggregates in YCFA only, YCFA + 1% DMSO, and the transition to a surface-coating biofilm across increasing concentrations of LCA. Scale bars = 500 µm.

Given that only LCA stimulated biofilm formation by EcN at concentrations that are physiologically relevant for a typical human colon, we focused on this bile acid for further investigation. We first asked whether oxygen tension affects LCA-induced biofilm formation in EcN, since a radial oxygen gradient is present in the colon.^29^ Intriguingly, modulating the ambient oxygen concentration revealed that EcN can form at least two distinct types of biofilms in YCFA, one of which requires oxygen but not LCA, while the other is responsive to LCA but not to oxygen. Under aerobic (20% O_2_) and microaerobic (1% O_2_) but not strictly anaerobic conditions, EcN formed a ring of biofilm at the air-liquid interface, which was enhanced under the microaerobic condition but formed regardless of the presence of LCA (Fig. 2C). By contrast, LCA triggered the formation of an extensive biofilm coating the bottom and sides of the culture vessel, regardless of the oxygen concentration (Fig. 2C).

To gain further insight into the nature of LCA-induced EcN biofilm formation, we used microscopy to examine the behavior of EcN in YCFA with and without LCA. In the absence of the bile acid, EcN formed aggregates during stationary phase in YCFA, both with and without the addition of 1% DMSO as a solvent control, although the aggregates formed in 1% DMSO appeared denser (Fig. 2D). However, as indicated by the crystal violet staining results for YCFA + 1% DMSO (Fig. 2A-B), these aggregates were non-adherent to the surface of the culture vessel. Addition of increasing doses of LCA induced a transition to a surface-coating biofilm in a graduated manner, whereby at the lowest biofilm-inducing concentration (25 µM), the aggregates started adhering to the bottom of the well, forming a web-like network, followed by progressive filling in of the gaps between the aggregates at higher concentrations (Fig. 2D).

### The adhesion of LCA-induced EcN biofilms is primarily protein-mediated

To identify which types of macromolecules comprise the matrix of the LCA-induced biofilm, we applied a combination of fluorescent dyes, enzymatic treatments, and mutagenesis. We first looked for extracellular DNA (eDNA). Staining 24-hr-old LCA-induced EcN biofilms with the cell-impermeant DNA dye TOTO-1 suggested little evidence of eDNA in the matrix, with the signal being localized to scattered spots likely representing dead cells (Fig. 3A).

**Fig. 3:**
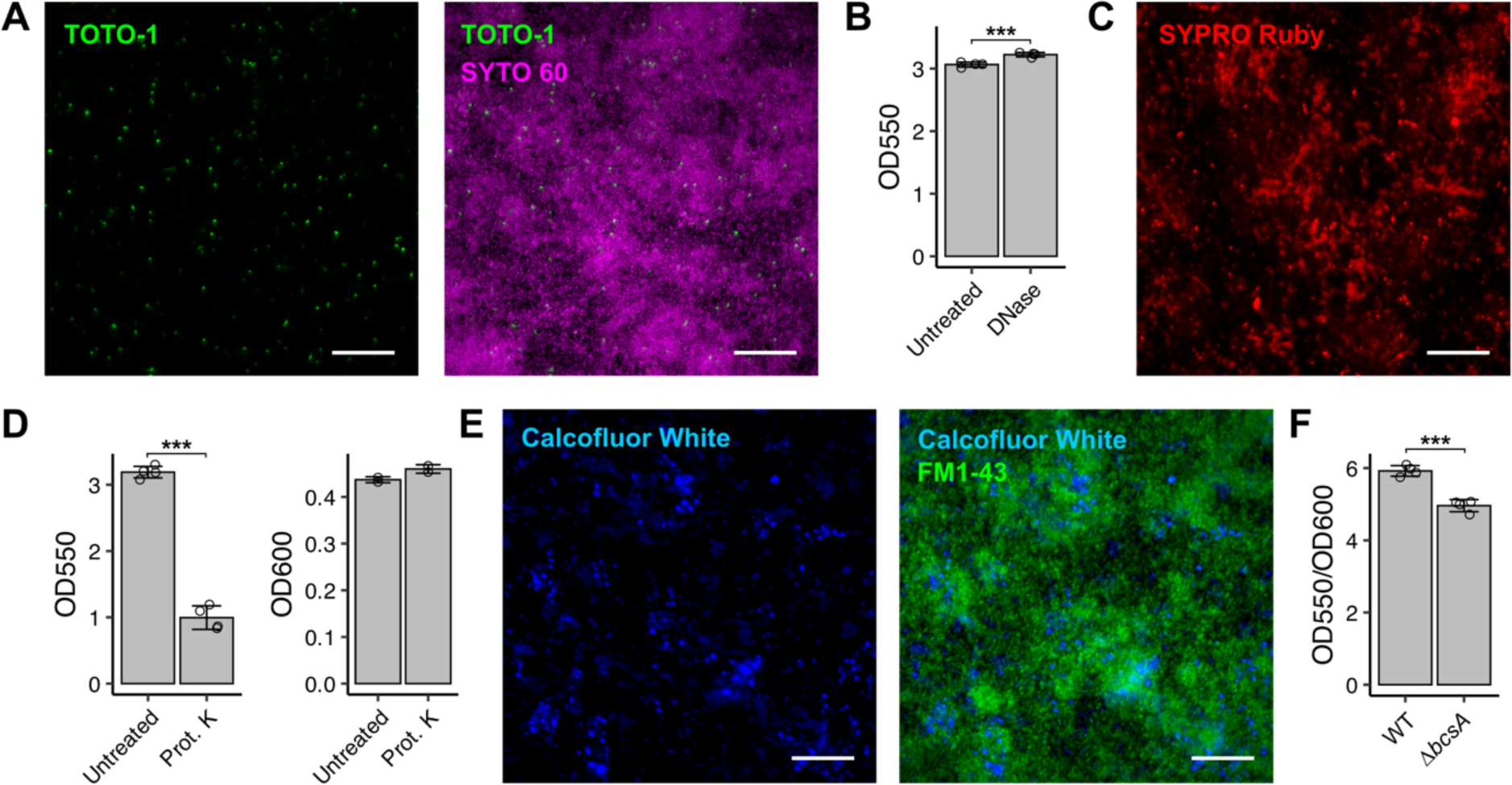
(A) Representative maximum intensity projection of an LCA-induced biofilm stained with TOTO-1 (extracellular DNA and dead cells) and SYTO 60 (intact cells). Scale bars = 20 µm. (B) Crystal violet quantification of biofilm biomass for cultures that were grown in the presence of LCA only (“untreated”) or LCA + DNase I for 24 hrs. *** *p* < 0.001, Welch’s t-test (*n* = 4). (C) Representative maximum intensity projection of an LCA-induced biofilm stained with SYPRO Ruby (general protein stain). (D) Crystal violet quantification of biofilm biomass, as well as OD_600_ measurement of total biomass (biofilm + planktonic), for cultures that were grown in the presence of LCA only (“untreated”) or LCA + proteinase K for 24 hrs. *** *p* < 0.001, Welch’s t-test (*n* = 4). (E) Representative maximum intensity projection of an LCA-induced biofilm stained with Calcofluor White (cellulose) and FM1-43 (cell membrane). Scale bars = 20 µm. (G) Quantification of biofilm formation (crystal violet absorption at OD_550_ normalized to OD_600_ as a proxy for growth) for the WT vs. Δ*bcsA* (cellulose-null mutant) after 24 hrs of growth in the presence of 100 µM LCA. *** *p* < 0.001, Welch’s t-test (*n* = 4).

Correspondingly, incubating the cultures with DNase I did not decrease the formation of LCA-induced biofilm; in fact, there was a very small but statistically significant increase in crystal violet staining in LCA-treated cultures grown in the presence of DNase I, compared to cultures grown without DNase (Fig. 3B). Imaging of LCA-induced biofilms labeled with the general protein stain SYPRO Ruby, on the other hand, suggested that proteins were present throughout the matrix (Fig. 3C). Indeed, incubating EcN with proteinase K severely hindered the formation of LCA-induced biofilms, without affecting overall growth (Fig. 3D). Finally, we stained LCA-induced biofilms with Calcofluor White to label cellulose, one of the major exopolysaccharides in the biofilms formed by many *E. coli* strains.^30^ The Calcofluor White signal appeared patchy (Fig. 3E), suggesting that EcN produces cellulose in the LCA-induced biofilm but that it is not homogenously distributed. To test the role of cellulose in LCA-induced biofilm formation, we knocked out *bcsA*, one of the genes involved in cellulose production. The mutant displayed a statistically significant decrease in crystal violet staining compared to the parent strain (Fig. 3F), but the magnitude of this decrease was much less than for the proteinase K treatment. Taken together, these results suggest that one or more proteinaceous factors serve as the major adhesin in LCA-induced EcN biofilms, with cellulose making a lesser contribution to the biofilm integrity and/or biomass.

### The mechanism of LCA-induced biofilm formation in EcN is multifactorial and distinct from air-liquid interface biofilms

To investigate the mechanism of LCA-induced biofilm formation in EcN at the molecular level, we first individually knocked out several adhesins and extracellular matrix components with known roles in *E. coli* biofilm formation or maturation, namely type I fimbriae (*fim*),^31,32^ F1C fimbriae (*foc*),^13^ the *E. coli* common pilus (*ecp*),^33^ curli fimbriae (*csgBA*),^34^ antigen 43 (*flu*),^35^ cellulose (*bcsA*, as previously mentioned),^36^ colanic acid (*wcaLM*),^37^ poly-*N*-acetylglucosamine (*pga*),^38^ and flagella (*fliC*).^39,40^ Consistent with the previously reported contributions of these genes to *E. coli* biofilm formation, each of these mutations caused a defect in the formation of the biofilm ring at the air-liquid interface, as indicated by a decrease of at least 50% relative to WT in crystal violet staining in the YCFA + 1% DMSO condition under low oxygen (Fig. 4A). However, only the loss of flagella (Δ*fliC*) caused a major reduction in LCA-induced biofilm, with the other mutations causing either no change or only mild defects up to a ∼25% reduction in crystal violet staining relative to the WT strain (Fig. 4A). These results suggested that LCA-induced biofilm formation proceeds via mechanisms that are at least partially distinct from formation of the biofilm ring at the air-liquid interface. In addition, while flagella are clearly important for both types of biofilms, LCA-induced biofilm formation must also involve at least one flagella-independent mechanism, as the *ΔfliC* mutant still displayed higher crystal violet staining in the presence of LCA relative to the DMSO control (Fig. 4B).

**Fig. 4:**
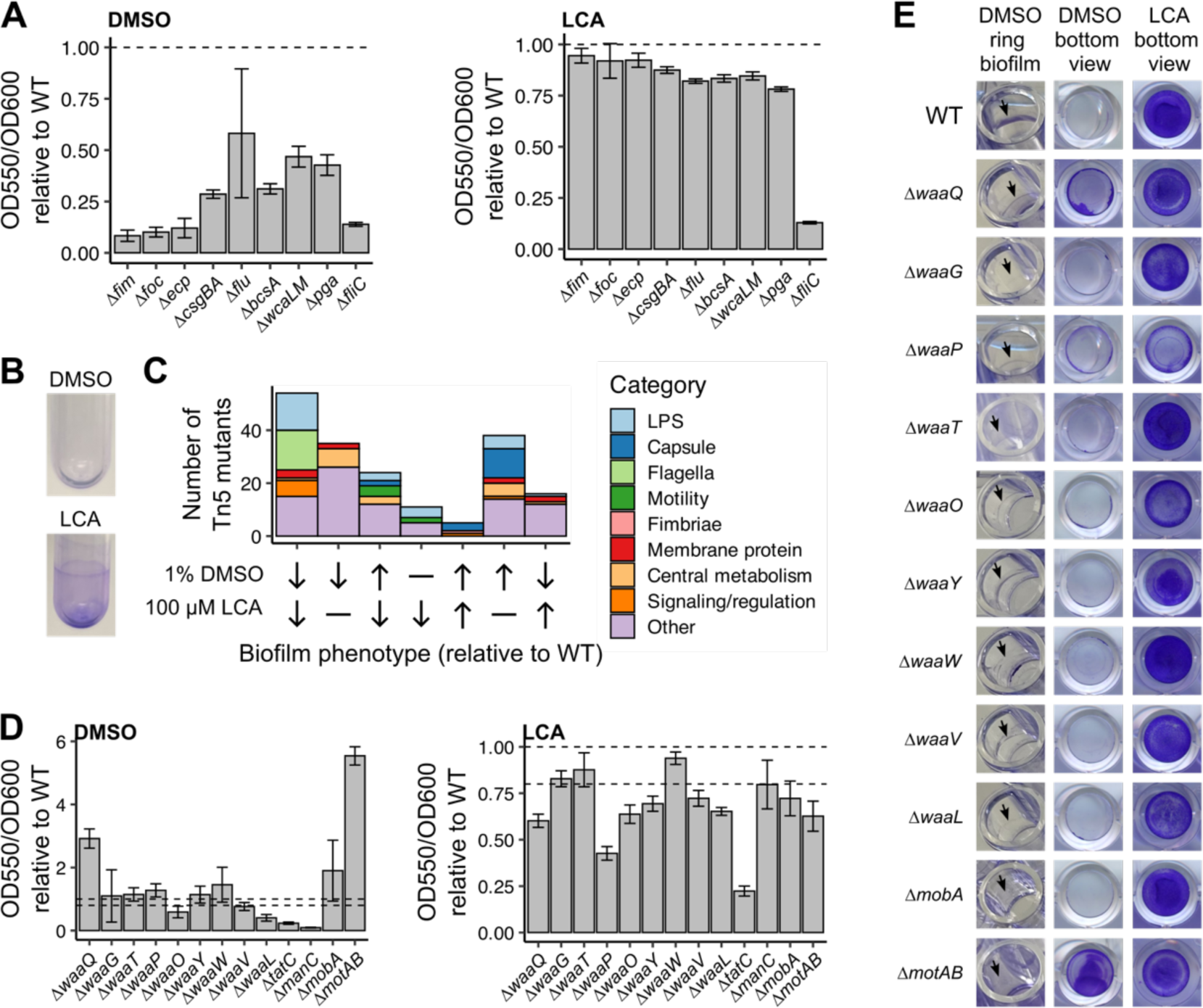
(A) Crystal violet quantification of relative biofilm formation (OD_550_/OD_600_ ratio normalized to that of the WT strain in the same condition) of targeted knockout strains grown in YCFA with 1% DMSO (solvent control) or 100 µM LCA. Error bars mark the standard deviation (*n* = 4 replicate wells from a representative experiment). (B) Images of the 5 mL tube crystal violet assay for Δ*fliC* grown microaerobically in YCFA with 1% DMSO vs. 100 µM LCA. (C) Summary plot of Tn5 screen showing the number of insertions per gene category and phenotypic effect on biofilm formation in 1% DMSO vs. 100 µM LCA. (D) Crystal violet quantification of relative biofilm formation (OD_550_/OD_600_ ratio normalized to that of the WT strain in the same condition) for lambda red knockouts of selected genes from the Tn5 screen. The upper dashed lines mark 100% of the WT biofilm level and the lower dashed lines mark 80% of the WT biofilm level. (E) Representative images of the 96-well plate crystal violet assay for the LPS gene knockouts, Δ*motAB*, Δ*mobA*, and the WT control, from the same experiment quantified in (D). The ring biofilm formed at the air-liquid interface in the DMSO control condition was visualized by photographing the plates at an angle (leftmost column). Black arrows mark the location of the air-liquid interface.

To identify other genetic factors involved in LCA-induced biofilm formation, we performed a random Tn5 transposon mutagenesis screen. In order to distinguish between mutations with a general impact on biofilm formation versus an LCA-specific impact, we screened the mutants in both YCFA + 1% DMSO and YCFA + 100 µM LCA, using the crystal violet microtiter plate assay as the readout. From a total of 10,066 colonies, we identified 184 unique mutants (1.8%) with a >20% decrease or increase in crystal violet staining in one or both conditions relative to the parent strain in the same condition, after normalizing to OD_600_ as a proxy for growth (Supplementary Data 1). Out of these, 53 mutants had a decrease in biofilm in both conditions, while 5 mutants had increased biofilm in both conditions. The other mutations either affected crystal violet staining in only one of the conditions, or caused opposite effects in the two conditions (Fig. 4C).

Among the mutants with a defect in LCA-induced biofilm, the most-represented functional category was genes related to LPS biosynthesis (21 unique insertions across 9 genes), followed by flagellar genes (17 unique insertions across 13 genes) (Fig. 4C, Supplementary Data 1). Interestingly, almost all mutants with a defect in LCA-induced biofilm also had disrupted or altered self-aggregation in YCFA + DMSO, suggesting that the mechanisms underlying these two phenomena may overlap to a large degree. None of the transposon insertions causing a defect in LCA-induced biofilm were in an annotated regulatory or signaling pathway except for *cyaA* (adenylate cyclase) (Supplementary Data 1), which is known to be required for expression of flagella.^41^

Similar to the phenotype of Δ*fliC*, the flagellar and *cyaA* transposon mutants were defective not only for formation of LCA-induced biofilm but also the air-liquid interface ring biofilm, as indicated by decreased crystal violet staining in the DMSO control condition (Supplementary Data 1). On the other hand, the LPS biosynthesis mutants displayed a variety of phenotypes, with some having decreased crystal violet staining in both conditions, while others had an LCA-specific decrease in staining, or even had increased staining in the DMSO control (Fig. 4C, Supplementary Data 1). The only other mutants with an LCA-specific decrease in crystal violet staining had insertions in *purA*, *opgH*, *tatC*, *manC*, *mobA*, or upstream of *lrhA* (Supplementary Data 1). However, the *purA* mutant also had a severe growth defect irrespective of LCA treatment. In addition, a second independent insertion in *opgH* caused decreased crystal violet staining in the DMSO control, suggesting that the role of this gene is not specific to LCA-induced biofilm (Supplementary Data 1). We therefore focused on *tatC*, *manC*, *mobA*, and *lrhA*, along with the *waa* locus responsible for biosynthesis of the LPS inner and outer core, for additional investigation. We also included the motility genes *motAB* because in contrast to mutations in the other structural components of flagella, transposon insertions in this locus caused an increase in crystal violet staining in the DMSO control, along with a moderate decrease in LCA-induced biofilm (Supplementary Data 1).

To verify that the transposon mutant phenotypes were due to loss of function of these genes (with the possible exception of *lrhA* given the insertion locations in a non-coding region), we generated in-frame deletions using lambda red recombineering and assessed the resulting phenotypes. Based on quantification of crystal violet staining, the LPS mutants Δ*waaP* and Δ*waaY* initially appeared to have a potentially LCA-specific defect, with Δ*waaP* having the most severe loss of LCA-induced biofilm among all the LPS core biosynthesis mutants (Fig. 4D). WaaP and WaaY catalyze the phosphorylation of heptose I and heptose II, respectively, in the LPS inner core.^42^ However, inspection of the staining pattern revealed that both mutants were in fact defective for formation of biofilm at the air-liquid interface, forming a thinner ring than the parent strain (Fig. 4E). The other LPS mutants either had minimal (<20%) decreases in LCA-induced biofilm (Δ*waaG*, Δ*waaT*, Δ*waaW*), similar or greater decreases in the DMSO control condition (Δ*waaO, ΔwaaV, ΔwaaL*), or a combination of decreased LCA-induced biofilm and increased crystal violet staining in the DMSO control condition (Δ*waaQ*), where the latter was due to increased adhesion on the bottom of the wells, rather than an increase in the air-liquid interface biofilm (Fig. 4E). Most of the LPS biosynthesis mutants were also defective for self-aggregation in YCFA + DMSO (Supplementary Fig. 2A), and all had reduced swimming motility to various degrees (Supplementary Fig. 2B).

Among the other deletion mutants, Δ*tatC*, Δ*manC*, and Δ*mobA* proved to have non-specific defects in biofilm formation (Fig. 4D-E). The phenotype of Δ*mobA* was potentially related to loss of the ability to respire DMSO—*mobA* is involved in biosynthesis of the molybdenum cofactor used by DMSO reductase,^43^ and the relatively loose, fluffy aggregates of Δ*mobA* in YCFA + DMSO resembled those formed by the parent strain in YCFA without DMSO (Supplementary Fig. 2A). On the other hand, Δ*motAB* formed a biofilm on the bottom of the wells in both YCFA + DMSO and YCFA + LCA, although neither biofilm was as extensive as the one formed by the parent strain in LCA (Fig. 4D-E). Despite forming such biofilms, this strain, which should possess paralyzed but otherwise structurally-intact flagella, failed to form the ring-like biofilm at the air-liquid interface (Fig. 4E), in addition to failing to self-aggregate in YCFA + DMSO (Supplementary Fig. 2A). Finally, for *lrhA*, which is a transcriptional repressor of flagella biosynthesis and motility,^44^ all three transposon insertions with a phenotype in our screen were upstream of the coding sequence, suggesting that the insertions could have caused overexpression of this gene rather than loss of function. RT-qPCR analysis confirmed that this was the case (Supplementary Fig. 2C). As expected, these transposon mutants also had decreased ability to swim in soft agar (Supplementary Fig. 2B), suggesting that their defects in biofilm formation were related to reduced motility and/or expression of flagella.

Taken together, the results of the transposon mutagenesis screen revealed that flagella play an important role during formation of LCA-induced biofilm that extends beyond enabling swimming motility. Alterations to the LPS core structure also affected LCA-induced biofilm formation, albeit in a pleiotropic manner, while the Δ*mobA* phenotype suggested a potential connection between anaerobic respiration, self-aggregation, and biofilm formation. However, no gene had a truly LCA-specific phenotype, as all transposon and deletion mutants with reduced crystal violet staining in YCFA + LCA also proved to have decreased formation of the ring-like air-liquid interface biofilm and/or altered self-aggregation in the solvent control condition.

### Flagellar components are upregulated by LCA and contribute to the biofilm extracellular matrix

Given the lack of transposon mutants with an LCA-specific defect, as well as the robustness of LCA-induced biofilm formation to the loss of several individual *E. coli* biofilm factors, we next asked whether LCA might influence biofilm formation through functionally-redundant pathways, which would not be detected in a screen based on single-locus mutants. To address this possibility, we first sought to characterize the transcriptional response of EcN to LCA. To this end, we performed an RNA-sequencing (RNA-seq) time course in which log-phase EcN cultures grown in anaerobic YCFA were exposed to 100 µM LCA, 100 µM CDCA, or 1% DMSO (the solvent control) for 15, 30, 90, and 180 minutes (Fig. 5A). CDCA was included as a control for genes that might respond to bile acids in general without being related to biofilm formation.

**Fig. 5:**
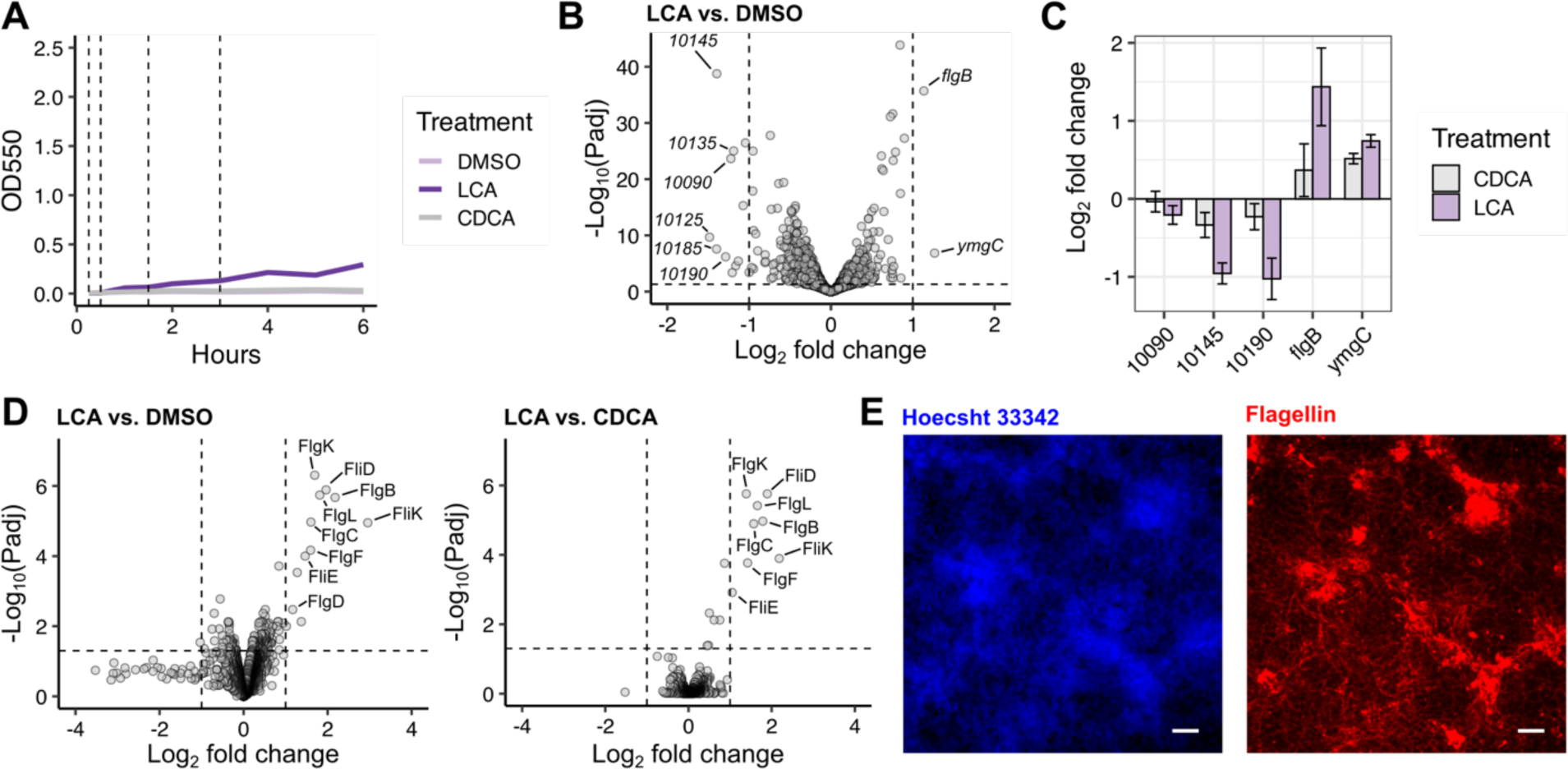
(A) Time course of biofilm accumulation under the conditions used for RNA-seq and proteomics (12-well plate cultures with starting OD_600_ of 0.5). Dashed vertical lines indicate the time points collected for RNA-seq (15, 30, 90, and 180 min). The y-axis is scaled to represent the values in relation to the typical OD_550_ (>2.0) of a 24-hr-old LCA-induced biofilm. (B) Volcano plot for the LCA vs. DMSO RNA-seq for samples collected at 180 min. The dashed vertical lines mark a log_2_ fold-change cutoff of +/- 1, while the dashed horizontal line marks an adjusted *p*-value cutoff of 0.05. (C) RT-qPCR assessment of the expression of *ECOLIN_10090, ECOLIN_10145, ECOLIN_10190, flgB*, and *ymgC* in EcN treated with DMSO vs. LCA vs. CDCA for 180 min. Samples were collected independently from the RNA-seq experiment. Fold changes were calculated with respect to the DMSO-treated control. Error bars represent the standard deviation (*n* = 3 replicates). (D) Volcano plots for LCA vs. DMSO and LCA vs. CDCA proteomics for samples collected at 180 min. The dashed vertical lines mark a log_2_ fold-change cutoff of +/- 1, while the dashed horizontal line marks an adjusted *p*-value cutoff of 0.05. (E) Representative images of Hoechst 33342 (DNA stain marking all cells; left) and anti-flagellin (right) staining of LCA-induced biofilm. The same field of view is shown in both images. Scale bar = 10 µm.

Analysis of the 15-minute and 30-minute time points revealed a lack of an immediate transcriptional response to either bile acid (Supplementary Fig. 3A-B). At 90 minutes, a small number of genes were upregulated by both bile acids relative to DMSO, including a cold-shock protein, genes involved in acid stress response, and the PTS glucose-specific subunit IIBC, while numerous genes involved in the transport or metabolism of other carbon sources (including allose, maltose, galactarate, and ethanolamine) were downregulated (Supplementary Fig. 3C, Supplementary Data 2). However, only two genes were also differentially expressed in the presence of LCA relative to CDCA at this time point: *yhhT* (annotated as an inner membrane/transport protein belonging to the AI-2 exporter superfamily) and a putative transposase, both of which were upregulated (Supplementary Fig. 3D, Supplementary Data 2). Although the putative function of *yhhT* hinted at a possible relation to biofilm formation given that AI-2 is a quorum sensing signal,^45^ RT-qPCR analysis suggested that, contrary to the RNA-seq results, this gene is not in fact upregulated by LCA (Supplementary Fig. 3E). To resolve this discrepancy, we inspected the RNA-seq read alignments, which revealed that the seeming upregulation of *yhhT* may have been driven by reads mapping to a highly-expressed downstream sequence that overlapped the tail end of the coding sequence of *yhhT* (Supplementary Fig. 3F). This downstream sequence mapped to a partial sequence of 23S rRNA; thus, we did not pursue it further.

At 180 minutes, by which time the LCA-specific increase in biofilm was evident (Fig. 5A), only LCA, and not CDCA, induced transcriptional differences relative to DMSO. Two genes, *flgB* (a flagellar basal body rod protein) and *ymgC* (part of a locus previously associated with biofilm regulation and acid stress response),^46^ were upregulated by LCA, while several genes belonging to a putative phage locus were downregulated (Fig. 5B, Supplementary Data 2). When CDCA was used as the reference for comparison instead of DMSO, the list of differentially expressed genes shrank to four of the putative phage genes (Supplementary Fig. 3G). Nevertheless, we were able to confirm the LCA-specific upregulation of *flgB*, and to a lesser extent *ymgC*, at this time point by RT-qPCR (Fig. 5C). While the functional importance of *flgB* for EcN biofilm formation had already been confirmed through our transposon mutagenesis screen, *ymgC* did not come up in the screen (Supplementary Data 1). Therefore, to explicitly test whether this locus contributes to LCA-induced biofilm formation, we generated an in-frame deletion mutant for *ymgBC* (*ymgB* is co-transcribed with *ymgC*^47^ and was also mildly upregulated by LCA, albeit below our fold-change cutoff). We also generated deletion mutants lacking different halves of the downregulated putative phage locus (avoiding a bacterial gene on the opposite strand in the middle of this locus), to test whether any of these genes might act as a repressor of biofilm formation, with that repression potentially being relieved by LCA. Neither loss of *ymgBC* nor loss of the phage genes had a major effect on LCA-induced biofilm formation (Supplementary Fig. 3H).

Overall, the RNA-seq results did not reveal a clear connection between an LCA-specific transcriptional response and biofilm induction. Nevertheless, given the importance of flagella for EcN biofilm formation, we were intrigued by the observation that only LCA upregulated *flgB* relative to the DMSO control at 180 minutes, and moreover that only *flgB*, among all the flagellar genes, was differentially expressed at the transcriptional level. Besides transcriptional regulation, flagellar genes and other biofilm factors can also be post-transcriptionally regulated.^48^ We therefore performed quantitative proteomics with tandem mass tag (TMT) labeling to investigate whether LCA affects the expression of additional genes at the protein level. For this experiment, samples were collected under the same conditions as the RNA-seq 180-minute time point. The proteomics analysis revealed that in addition to FlgB, seven other flagellar proteins were upregulated by LCA relative not only to DMSO, but also to CDCA; moreover, these were the only differentially expressed proteins in the latter comparison (Fig. 5D). All of the upregulated genes belonged to the class II flagellar genes (i.e. those that are expressed in the middle of the temporal cascade of flagellar gene expression),^49^ suggesting that we may have caught the cells midway through the flagella production process at this time point.

In an attempt to further investigate the effect of LCA on flagellar expression, we performed scanning electron microscopy (SEM) to directly assess the presence and morphology of flagella in cultures treated with 100 µM LCA or 1% DMSO for 24 hours, focusing on the planktonic fractions due to sample preparation considerations and to provide a fairer basis for comparison. Unfortunately, the presence of aggregates and cell-to-cell heterogeneity made it difficult to determine whether there was any difference in flagellar expression between the two conditions. However, we noted that in both conditions, the cells in the aggregates appeared to be enmeshed in a network of flagella (Supplementary Fig. 4). We therefore stained LCA-induced biofilms grown on glass coverslips with an anti-flagellin antibody, which revealed a similar mesh-like network throughout the biofilm (Fig. 5E). Taken together, LCA upregulates the production of class II flagellar components at a timepoint that corresponds to early biofilm formation, potentially via both transcriptional and post-transcriptional mechanisms. Furthermore, flagella appear to represent a major component of the extracellular matrix of LCA-induced biofilm, as well as the non-surface-attached aggregates that EcN forms in YCFA in the absence of LCA.

### LCA-induced biofilm formation impacts antibiotic tolerance and inter-strain competition

We next asked whether LCA-induced biofilm formation could confer any fitness benefits to EcN. We started by testing its effect on antibiotic tolerance, since increased antibiotic tolerance is a broadly-conserved characteristic of bacterial biofilms.^50,51^ We focused on ciprofloxacin because it is one of the most commonly-prescribed antibiotics for patients with inflammatory bowel disease,^52^ an indication for which EcN has shown therapeutic potential in clinical trials.^53^ After verifying that LCA did not affect the minimum inhibitory concentration (MIC) of ciprofloxacin, we grew EcN with 100 µM LCA for 24 hours to allow it to form a biofilm. We then separated the planktonic fraction from the biofilm fraction and treated both with high concentrations of ciprofloxacin for another 24 hours. In comparison to the LCA-treated planktonic cells, the LCA-induced biofilms were approximately 20-fold more tolerant to 0.5 µg/mL ciprofloxacin, which is 32x the MIC (Fig. 6A). However, when the planktonic and biofilm fractions were not separated, LCA did not increase the overall percentage of surviving cells compared to the DMSO control (Fig. 6B). This might suggest that the aggregates formed in 1% DMSO, though not surface-attached, also represent a biofilm-like physiological state with respect to increased antibiotic tolerance, similar to what has been observed in other bacterial species.^54^

**Fig. 6:**
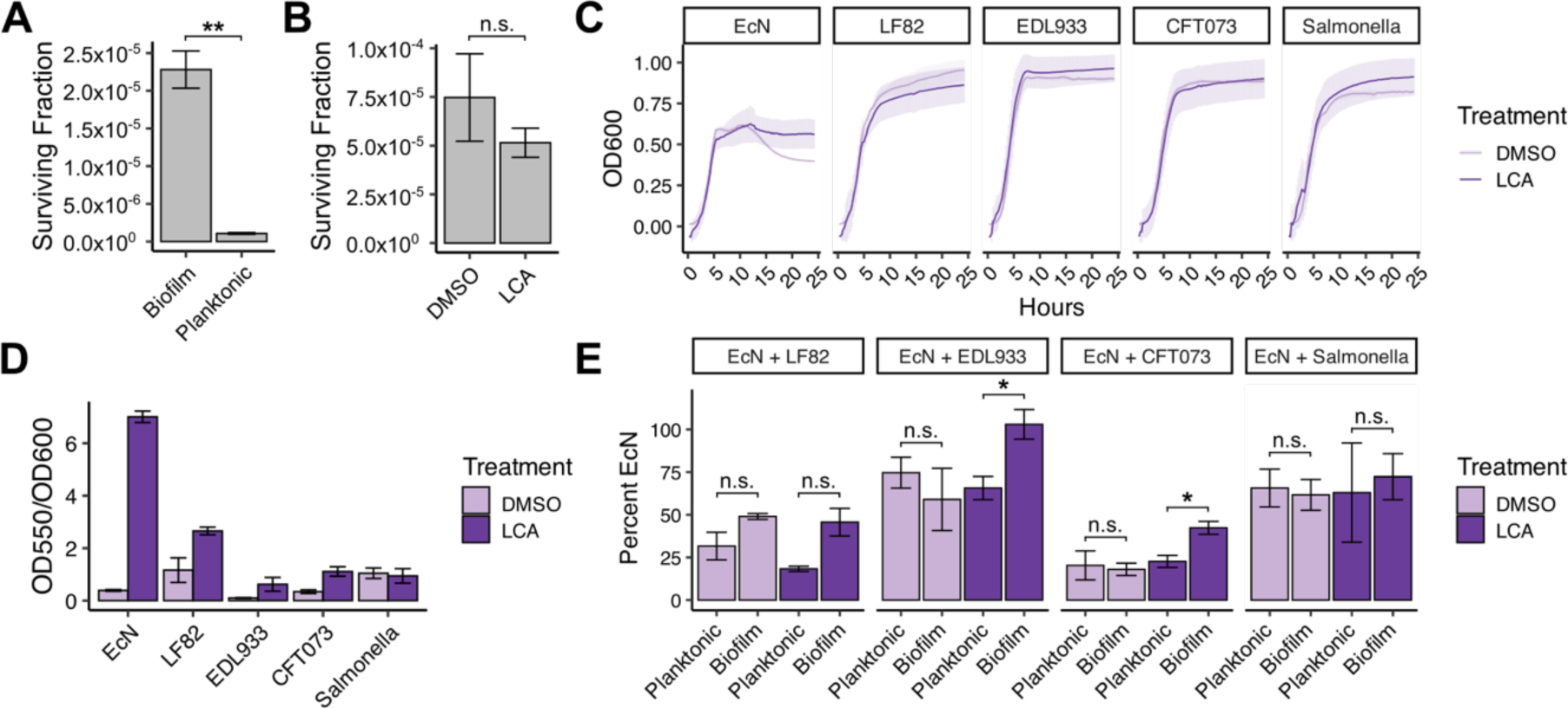
(A) The fraction of surviving cells after 24 hrs of treatment with 0.5 µg/mL ciprofloxacin, for LCA-induced biofilm vs. planktonic cells collected separately from the same cultures. Error bars represent the standard deviation (*n =* 3 replicates). ** *p* < 0.01, Welch’s t-test. (B) The fraction of surviving cells after 24 hrs of treatment with 0.5 µg/mL ciprofloxacin, for LCA-treated vs. DMSO-treated cultures in which biofilm and planktonic cells were not collected separately. Error bars represent the standard deviation (*n =* 3 replicates); n.s. = not significant (*p* > 0.05, Welch’s t-test). (C) Growth curves in the presence of 1% DMSO vs. 100 µM LCA for EcN and the other enteric strains used in competition experiments. The drop in OD_600_ for EcN upon entering stationary phase in 1% DMSO likely reflects the onset of self-aggregation (see Fig. 1D). Error bars represent the standard deviation (*n* = 6 replicates). (D) Quantification of biofilm formation (crystal violet absorption at OD_550_ normalized to OD_600_) after 24 hrs of treatment with 1% DMSO or 100 µM LCA, for the strains used in competition experiments. Error bars mark the standard deviation (*n* = 4 replicate wells from a representative experiment). (E) The percentage of the planktonic or surface-attached (biofilm) final population achieved by EcN, based on CFU counts, after 24 hrs of growth in pairwise competition with other strains, in the presence of 1% DMSO or 100 µM LCA. Error bars represent the standard deviation (*n* = 3 replicates). * *p* < 0.05, biofilm vs. planktonic within each treatment (Welch’s t-test followed by the Bonferroni correction for multiple comparisons).

We also investigated whether LCA-induced biofilm formation could affect the outcome of competition between EcN and other enteric bacteria. For these experiments, we used the enteric pathogens *Salmonella enterica* subsp*. enterica* serovar Typhimurium strain ATCC 14028, enterohemorrhagic *E. coli* (EHEC) O157:H7 strain EDL933, and adherent invasive *E. coli* (AIEC) strain LF82, along with the uropathogenic strain *E. coli* CFT073, which is a close genomic relative of EcN.^7^ 100 µM LCA did not significantly affect the individual growth of these strains (Fig. 6C), nor did it induce biofilm formation in any of the strains to the same extent as for EcN (Fig. 6D). Following these preliminary tests, kanamycin-resistant EcN was inoculated at a 1:1 ratio with each competing strain in the presence of either 1% DMSO or 100 µM LCA. After 24 hours, the planktonic and biofilm fractions of the co-cultures were plated separately for CFUs. In co-cultures with AIEC LF82, EcN appeared to compete more successfully in the biofilm population compared to the planktonic phase even in the DMSO control, but this difference was not statistically significant, and its fitness was not further enhanced by LCA (Fig. 6E). By contrast, in co-cultures with EHEC EDL933, LCA significantly improved the competitive fitness of EcN, resulting in essentially complete exclusion of EHEC EDL933 from the biofilm population (Fig. 6E). LCA also helped EcN reach a higher proportion of the biofilm population in co-cultures with *E. coli* CFT073 (Fig. 6E). In competition with *Salmonella*, EcN reached a similar proportion of both the planktonic and biofilm populations, regardless of treatment with LCA (Fig. 6E). Overall, these results suggested that LCA can indeed enhance the competitive fitness of EcN specifically during biofilm formation, although this effect is not universal across all competing strains.

### LCA promotes EcN biofilm formation on human colonic epithelial monolayers

Finally, we asked whether LCA could increase EcN adherence and biofilm formation not only on abiotic surfaces, but also on a model of the human intestinal epithelium. To test this, we co-cultured GFP-tagged EcN with epithelial monolayers that were derived from human colon organoids and differentiated on Transwell supports (Fig. 7A). Organoid-derived monolayers have previously been shown to recapitulate key aspects of the *in vivo* intestinal epithelium, including the presence of a soluble mucus layer,^55,56^ making them a useful model for studying host-microbe interactions in the human gut. Mucus-producing goblet cells were abundant in our differentiated monolayers (Fig. 7B), and we noticed that the apical media became highly viscous during the course of our experiments, suggesting ample mucus production. Consistent with prior studies performed with a colorectal cancer-derived cell line,^57,58^ EcN was able to adhere to the epithelial monolayers to some extent and form microcolonies even in the absence of any bile acid (Fig. 7C). However, the addition of 100 µM LCA significantly increased the overall level of adherence and led to the formation of large mats of EcN on the monolayers, which were resistant to multiple washes (Fig. 7C-D). 100 µM CDCA, which we used as a non-biofilm-inducing control bile acid condition, did not produce the same effect (Fig. 7C-D). Neither the bile acids, the co-culture media, nor EcN itself demonstrated toxicity towards the epithelial cells as measured by a lactate dehydrogenase (LDH) release assay (Supplementary Fig. 5), although we did note a minor increase in toxicity for the combination of EcN and LCA. This mild synergistic toxicity may be an artifact of our artificial closed culture system, given that EcN has repeatedly proved to be safe in clinical trials.^6^ Taken together, these results suggest that our experiments on abiotic surfaces were predictive of EcN’s ability to form biofilms on colonic epithelial cells, and that LCA could potentially exert a similar effect *in vivo*.

**Fig. 7:**
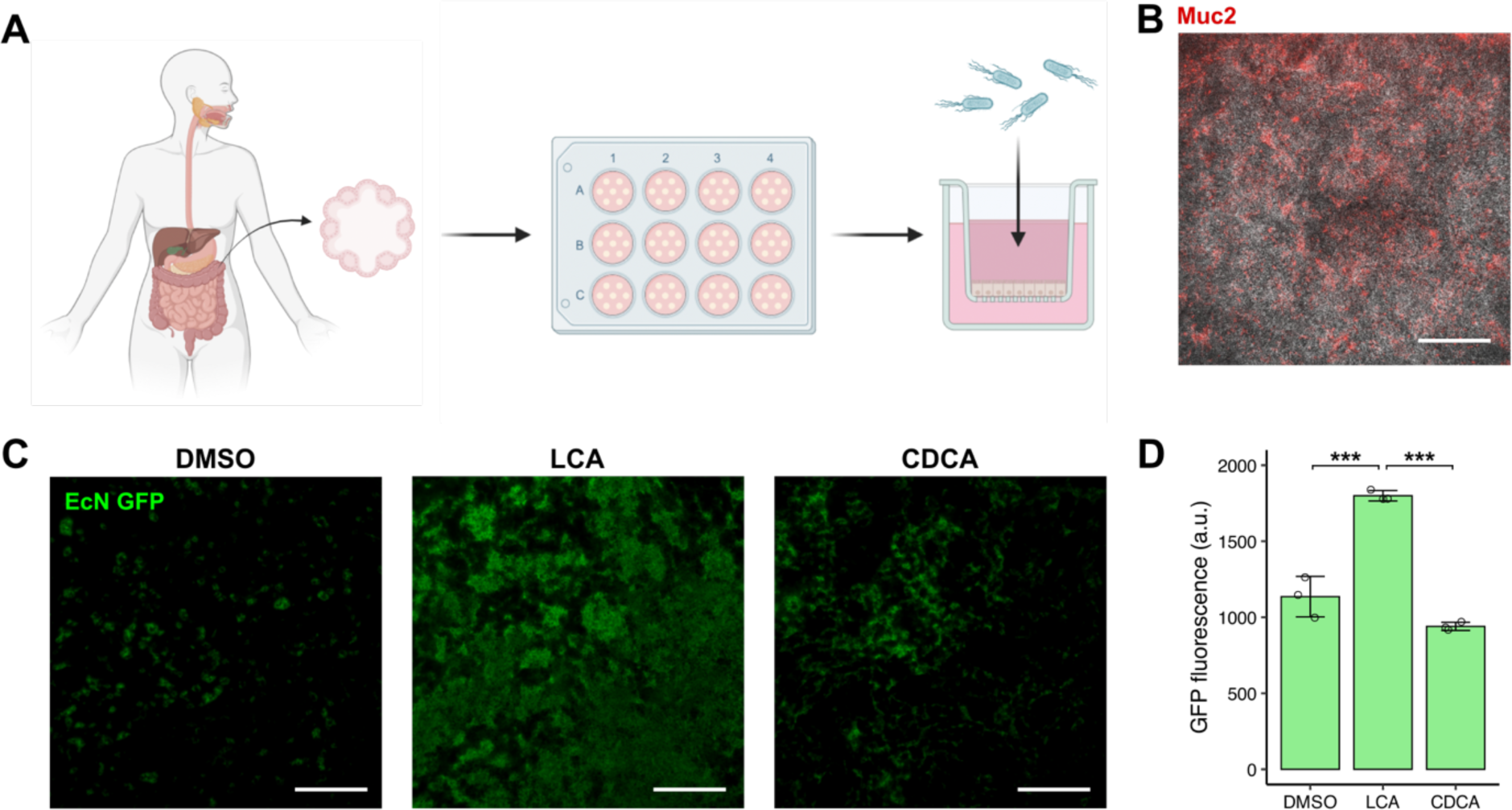
(A) Experimental schematic for testing the effect of LCA on EcN adhesion to the intestinal epithelium. Colonic organoids were derived from intestinal adult stem cells and expanded in 3D culture in multi-well plates before being dissociated and seeded on Transwell membranes to form monolayers. Following differentiation of the monolayers, EcN was added to the apical chamber. The schematic was created in Biorender. (B) A representative organoid monolayer stained for intracellular Muc2 (red) to mark goblet cells, overlaid on a phase contrast image, following 5 days of differentiation. Scale bar = 500 µm. (C) Representative images of GFP-expressing EcN adhering to organoid monolayers after 8 hrs of co-culture in the presence of 1% DMSO, 100 µM LCA, or 100 µM CDCA. The apical surface of the monolayers was washed three times with PBS prior to imaging to remove non-adherent bacteria. Scale bars = 500 µm. (D) Quantification of the GFP signal from the experiment shown in (C) following lysis and homogenization of each well. *** *p* < 0.001 (ANOVA followed by Tukey HSD).

## DISCUSSION

Here, we have shown that the gut-microbiota-derived secondary bile acid LCA can induce the formation of a distinctive surface-coating biofilm by the probiotic strain *E. coli* Nissle 1917. The human gut environment is heterogenous and dynamic, and biofilm formation by EcN *in vivo* is likely subject to variations in bile acid concentrations, nutrient availability, gut microbiota composition, and other factors. Nevertheless, several observations suggest that LCA could influence how EcN colonizes the human gut. First, LCA is typically one of the most abundant bile acids in the human colon, and in our assays, it promoted biofilm formation by EcN at concentrations within the ranges reported for human feces and cecal contents. Second, formation of this surface-coating biofilm remained robust when LCA was mixed with other bile acids. Third, LCA-induced biofilm formation occurred in a medium that is effective for culturing a broad range of gut bacteria *in vitro*,^26^ suggesting that it captures major features of the nutritional profile of typical human colon contents. Fourth, formation of this type of biofilm was insensitive to oxygen concentration, which varies within the colon depending on distance from the mucosal surface,^29^ as well as during times of inflammation versus homeostasis.^59^ Finally, LCA triggered increased adhesion of EcN not only on abiotic surfaces, but also on mucus-producing epithelial monolayers derived from human colon organoids.

Bile-acid-induced biofilm formation has previously been reported in several other species of gut bacteria, including *Clostridioides difficile*, *Bifidobacterium*, *Lactobacillus*, *Vibrio cholerae*, *Bacteroides fragilis*, *Bacteroides thetaiotaomicron*, *S. enterica*, *Shigella flexneri,* and *Enterococcus faecalis*.^16–19,23,60–63^ Many of these studies were performed with bile extract or mixed bile acids rather than with individual bile acids. Taken together, these reports suggest that bile components are widely used by gut bacteria as signals for triggering biofilm formation. However, the studies based on purified bile acids have revealed that a given species usually responds preferentially to only a subset of bile acids, and different species and strains can respond differently to the same bile acid.^17,23,64,65^ Our findings support these trends, as EcN responded most dramatically to LCA among the nine bile acids we tested, and the other strains of *E. coli* that we tested did not show a similar magnitude of response. Interestingly, to our knowledge, *E. faecalis* and *B. thetaiotaomicron* are the only other species reported so far to have strains that respond preferentially to LCA in terms of biofilm formation.^23,64^ Several other species reported to form bile-acid-induced biofilm instead respond to primary bile acids conjugated to glycine or taurine, as in the case of *Bifidobacterium*, *Lactobacillus*, and *Salmonella*,^16,60,65^ while *C. difficile* forms biofilms in response to both the deconjugated primary bile acid CDCA and the secondary bile acid DCA, but not LCA.^17^ Conjugated primary bile acids are the main constituents of bile in the small intestine, whereas deconjugated primary and secondary bile acids dominate in the cecum and colon.^66,67^ Thus, the segment of the gastrointestinal tract where bile-acid-induced biofilm formation is relevant likely differs for different species.

Several different mechanisms have been proposed for bile-acid-induced biofilm formation, again depending on the species in question. These include increased production of extracellular polysaccharides,^62^ cyclic-di-GMP,^65^ autotransporter adhesins,^68^ curli fimbriae,^69^ or eDNA.^17^ Additionally, an endogenous extracellular DNase is required for maximal bile-induced biofilm formation in *B. thetaiotaomicron*,^19^ while LCA-induced biofilm formation in *E. faecalis* is associated with a morphotype switch from diplococci to long chains of cells.^23^ However, the mechanism by which LCA induces surface-coating biofilm formation in EcN likely differs from all of the above. We found no evidence of LCA-induced upregulation of extracellular polysaccharides, autotransporters, curli, or other fimbriae in EcN, either by transcriptomics or proteomics. Moreover, eDNA was only minimally present in LCA-induced biofilms, which were insensitive to DNase treatment, and we did not observe cell chaining.

What, then, could be the mechanism(s) driving LCA-induced biofilm formation in EcN? While we did not find evidence of a gene or signaling pathway that regulates an LCA-specific response, the LCA-induced biofilm phenomenon may be related in part to the inherent tendency of EcN to self-aggregate during stationary phase in YCFA, as nearly all biofilm-deficient mutants in our transposon mutagenesis screen also displayed abnormal or reduced self-aggregation. Indeed, phase-contrast images of EcN’s response to the lowest biofilm-promoting dose of LCA (25 µM) suggested that rather than disrupting or preventing self-aggregation of the bacteria, LCA likely promotes the adhesion of the aggregates to the surface. With increasing concentrations of LCA, the adhesion of the aggregates appeared to be combined with either spreading along the surface or increased adhesion of individual bacteria to fill the spaces in between the aggregates.

This shift from floating aggregates to a surface-coating biofilm could be driven in part by the increased production of flagellar components during early biofilm formation in the presence of LCA, as flagella can promote adhesion to hydrophobic surfaces.^70^ The polystyrene microtiter plates used in most of our experiments are hydrophobic, and an adhesive role for flagella during LCA-induced biofilm formation is further suggested by the fact that mutants lacking structural components of flagella had a much more dramatic defect compared to mutants with paralyzed flagella. The gastrointestinal mucosal surface is also thought to be hydrophobic, with a gradient of increasing hydrophobicity from the duodenum to the distal colon.^71^ Thus, flagella-mediated adhesion may also be relevant *in vivo*. Indeed, our SEM images of EcN aggregates, as well as the staining pattern of flagellin in LCA-induced biofilms, are reminiscent of previously published micrographs of EcN adhering to cultured porcine intestinal epithelial cells, in which the bacterial cells were similarly surrounded by a mesh-like network of flagella.^72^ A flagella-null mutant also showed reduced adhesion in that study. Taken together, these findings suggest that self-aggregation and surface adhesion via flagella may help EcN colonize the intestinal mucosa, and that this behavior could be enhanced by LCA. LCA itself might also directly interact with flagella, as it is highly hydrophobic. However, it remains unclear how or why the expression of flagellar components is upregulated in the presence of LCA. Moreover, as previously noted, there must also be a flagella-independent mechanism by which LCA promotes biofilm formation, since loss of flagella did not completely negate the increased surface adhesion of EcN in the presence of LCA.

Given that flagella were the only LCA-specific upregulated cellular component at the protein level during early biofilm formation, the flagella-independent mechanism of LCA-induced adhesion may involve a physical interaction between LCA and pre-existing features of the EcN cell surface. The results of our transposon mutagenesis screen suggest two primary candidates: LPS and the K5 capsule. Multiple mutations affecting the LPS core structure caused a reduction in LCA-induced biofilm despite increasing the propensity of EcN to stick to the bottom edges of the well in the absence of LCA. However, these reductions in LCA-induced biofilm could be related to the reduced motility of the LPS mutants or other indirect effects, rather than an altered interaction between LCA and LPS. Indeed, a direct interaction between LCA and LPS would be somewhat surprising, since the O-antigen and core oligosaccharide of LPS are hydrophilic, whereas LCA is highly hydrophobic, and both LPS and LCA are negatively charged at physiological pH.^73,74^ A potential interaction between LCA and the K5 capsule, on the other hand, is suggested by the fact that most of the transposon mutants with a strong surface-coating biofilm in the absence of LCA had insertions in genes for capsule biosynthesis—consistent with a previous report of biofilm gain-of-function for capsule-deficient mutants.^58^ Thus, if LCA binds to the capsule and either disrupts it or increases its hydrophobicity, this could lead to the increased surface adhesion of EcN. However, like LPS, the K5 capsule is hydrophilic and negatively-charged,^75^ which might make such an interaction unlikely. If not LPS or the capsule, the lack of other clear candidates from the transposon mutagenesis screen suggests that LCA might bind non-specifically or have an affinity for multiple functionally-redundant surface features, which would be challenging to detect via traditional genetic approaches.

Regardless of the specific mechanism by which LCA stimulates the formation of a surface-coating biofilm, the results of our functional assays suggest the intriguing possibility that LCA-induced biofilm formation may in some situations contribute to the probiotic properties of EcN. Pathogen suppression is thought to be one of the beneficial properties of EcN,^6^ and LCA helped EcN better compete against two out of four tested pathogenic strains during surface colonization. This is consistent with an earlier report that EcN demonstrated a higher competitive fitness against various intestinal pathogens during biofilm formation compared to planktonic growth.^14^ However, whereas those earlier experiments were conducted in a minimal medium that is not representative of the colon environment, our findings offer new insight into how biofilm formation and the competitive fitness of EcN might be regulated *in vivo*. EcN is also known to be able to suppress other strains irrespective of biofilm formation through the production of microcins, but this only occurs under iron-limited conditions, such as during inflammation.^11^ By contrast, LCA is present under homeostatic conditions in humans. Unfortunately, testing the specific impact of LCA-induced biofilm formation on the competitive fitness and probiotic properties of EcN *in vivo*, e.g. in mice, would likely be challenging. Unlike in humans, LCA is normally only minimally present in rodents,^76^ and exogenous LCA feeding can have serious detrimental consequences, including hepatobiliary injury leading to eventual death.^77^ The Transwell setup we used for our organoid-derived monolayers also has limited utility for studies on probiotic effects, because extended co-cultures lead to bacterial overgrowth and cell death. However, the recent development of human “mini-colons” with long-term viability in a microfluidic platform might offer one avenue for follow-up studies on this topic.^78^

More broadly, our findings re-emphasize the importance of considering the natural environment of a bacterium when studying its behavior *in vitro*. Under conditions simulating some of the nutritional aspects of the colon environment, addition of a gut microbiota-derived bile acid triggered the formation of a physically and mechanistically distinctive biofilm by a widely-studied probiotic strain, which would otherwise be missed. Biofilm formation by gut bacteria is a topic of increasing interest not only in the context of probiotics, where biofilm formation might be desirable, but also because several studies have identified a correlation between various gastrointestinal disorders and the presence of mucosal biofilms detectable by microscopy or endoscopy.^79–83^ Moreover, the oncogenic potential of the bacterially-produced genotoxin colibactin depends on bacterial adhesion to the intestinal epithelium.^84^ Yet for many gut bacteria, biofilm formation and regulation remain poorly understood—and even among well-studied species, there can be significant strain-level differences, as we have shown here for EcN in comparison to pathogenic strains of *E. coli*. Indeed, in a recent study, the majority of isolates from biofilm-positive intestinal biopsies failed to form biofilms under standard culture conditions,^79^ suggesting that some of those strains may also require a gut-mimicking environment to trigger biofilm formation. Incorporating different bile acids and other gut-specific metabolites into future studies may shed light on biofilm regulation in a wider range of strains and species, with potential implications for functional engineering of the human gut microbiota.

## MATERIALS AND METHODS

### Strain construction

All strains, plasmids, and primers used in this study are listed in Supplementary Table 2. For most experiments in this study, we used *E. coli* Nissle 1917 (isolated from Mutaflor® and authenticated by whole genome sequencing) that had been cured of its native plasmids, pMut1 and pMut2, to facilitate further genetic manipulations. For experiments testing the impact of transposon insertions or gene deletions on biofilm formation, we used EcN Δ*clbQ*::*kan* (which lacks a gene in the colibactin biosynthesis pathway) as the “WT” reference strain to enable the use of kanamycin throughout the experiments. EcN Δ*clbQ*::*kan* was also used as the kanamycin-resistant strain in competition experiments with other enteric bacteria. The original strain isolated from Mutaflor®, the plasmid-cured strain, and EcN Δ*clbQ*::*kan* all behaved identically with respect to biofilm formation and the response to LCA.

To generate plasmid-cured EcN, linear pMut1 and pMut2 were first amplified by PCR using the native pMut1 and pMut2 plasmids isolated from EcN as a template. The *sacB* gene and ampicillin resistance cassette and *sacB* and gentamicin resistance cassette were amplified using pEX18gm and pEX100T as templates, respectively.^85,86^ The recombinant pMut1-sacB-gmR and pMut2-sacB-ampR plasmids were constructed using NEBuilder HiFi DNA Assembly Master Mix (New England Biolabs) according to the manufacturer’s instructions, and authenticated via whole plasmid sequencing. Subsequently, the recombinant suicide vectors were introduced into electrocompetent EcN. First, the pMut1-sacB-gmR suicide vector was selected for using gentamicin at a final concentration of 10 µg/mL in LB agar. Upon isolation of single colonies, the pMut1-sacB-gmR plasmid was counter selected for in the presence 10% sucrose in LB low salt agar medium to obtain an *E. coli* Nissle strain containing only the native pMut2 plasmid. The same method was then used to cure native pMut2 using the pMut2-sacB-ampR suicide vector, instead using ampicillin at a final concentration of 100 µg/mL in LB agar for selection of the recombinant suicide vector followed by counterselection on 10% LB low salt agar medium. PCR using primers previously designed to selectively amplify pMut1 and pMut2 was performed on minipreps of the plasmid-cured strain to confirm the absence of both cryptic native plasmids.^87^

Strains with gene deletions were constructed using standard lambda Red recombineering techniques. Briefly, plasmid-cured EcN was transformed with pSIM18 containing the *exo, bet,* and *gam* genes expressed from the *pL* operon under the control of a temperature-sensitive (ts) repressor, CI857, and a hygromycin resistance cassette.^88^ Transformants were selected on LB agar containing 200 µg/mL hygromycin at 30°C. The kanamycin resistance cassette was amplified from pKD4 by PCR using primers with 50 bp homology arms flanking the gene targeted for deletion.^89^ Meanwhile, a culture of EcN pSIM18 was grown to mid-log phase at 30°C in LB + 200 µg/mL hygromycin, then expression of the lambda Red proteins was induced by incubating the cells at 42°C for 15 min, followed by 15 min on ice. Electrocompetent cells were then prepared by two successive washes and concentration in ice-cold 10% glycerol. Following DpnI digestion and purification with the QIAquick PCR Purification Kit (Qiagen), the pKD4 PCR product was electroporated into EcN pSIM18, which was then recovered in 1 mL SOC medium (Invitrogen) at 37°C with shaking at 200 rpm for four hours. The transformation was plated on LB agar supplemented with 50 µg/mL kanamycin and incubated at 37°C overnight. Primers annealing internally and externally to the targeted gene and the kanamycin resistance cassette were used to screen single colonies for correct integrants. Confirmed deletion mutants were maintained under kanamycin selection during all experiments.

EcN pMut1-sfGFP was constructed by first PCR amplifying linear pMut1. A gBlock^TM^ containing the J23119 promoter (Registry of Standard Biological Parts BBa_J23119), superfolder GFP^90^ and chloramphenicol resistance cassette was synthesized by Integrated DNA Technologies (IDT). The final plasmid was constructed using NEBuilder HiFi DNA Assembly Master Mix (New England Biolabs) according to the manufacturer’s instructions and authenticated via whole plasmid sequencing. The plasmid was then electroporated into plasmid-cured EcN and transformants were selected on LB agar supplemented with 12.5 µg/mL chloramphenicol. Because pMut1 is stably maintained in EcN without selection, chloramphenicol was omitted once the strain had been isolated by restreaking.

### Culture media and growth conditions

EcN was routinely grown aerobically at 37°C in LB broth with shaking at 200 rpm. For growth on solid media, LB was supplemented with 15 g/L agar. Biofilm experiments were performed in YCFA media, consisting of the following ingredients per liter: tryptone (10 g), yeast extract (2.5 g), D-glucose (2 g), D-maltose (2 g), D-cellobiose (2 g), MgSO_4_ (0.09 g), CaCl_2_ (0.09 g), K_2_HPO_4_ (0.45 g), KH_2_PO_4_ (0.45 g), NaCl (0.9 g), (NH_4_)_2_SO_4_ (0.9 g), resazurin (1 mg), NaHCO_3_ (4 g), L-cysteine (1 g), hemin (10 mg), acetic acid (4.05 mL), propionic acid (1.43 mL), isobutyric acid (0.24 mL), *n*-valeric acid (0.24 mL), isovaleric acid (0.24 mL), biotin (0.01 mg), folic acid (0.05 mg), pyridoxine (0.15 mg), thiamine hydrochloride (0.05 mg), riboflavin (0.05 mg), cyanocobalamin (0.01 mg), and *p*-aminobenzoic acid (0.03 mg). The media was prepared with Milli-Q water and the vitamins were added separately from a 1000x filter-sterilized solution after autoclaving the rest of the media and cooling to 55°C. Following preparation of YCFA, the bottle caps were closed tightly and wrapped in Parafilm to minimize oxygen diffusion, and the bottles were stored under ambient conditions. Upon opening a new bottle of YCFA, the media was passively degassed inside an anaerobic chamber (Coy Laboratory Products) until the resazurin became colorless, and the media was subsequently stored inside the chamber. YCFA media was used within four months of the preparation date.

The following bile acids were used in this study: TCDCA (MedChemExpress, catalog no. HY-N2027), GCDCA (AstaTech, catalog no. F85524), CDCA (Sigma-Aldrich, catalog no. C9377), UDCA (Sigma-Aldrich, catalog no. U5127), LCA (Sigma-Aldrich, catalog no. L6250), TCA (MedChemExpress, catalog no. HY-B1788), GCA (Cayman Chemicals, catalog no. 20276), CA (Sigma-Aldrich, catalog no. C1129), and DCA (Sigma-Aldrich, catalog no. D2510). Bile acid powder stocks were stored according to the manufacturer’s instructions. 100x stock solutions were prepared by dissolving the respective bile acid in DMSO (Sigma-Aldrich, catalog no. D2650).

For anaerobic growth, experiments were typically set up under ambient conditions and subsequently incubated statically at 37°C inside an anaerobic chamber (Coy Laboratory Products), except for the RNA-seq time course, RT-qPCR, and proteomics experiments, which were not only incubated anaerobically but also set up under anaerobic conditions given the relatively short experimental timeframe. Multi-well plate assays that were performed anaerobically were incubated inside gallon-size plastic Ziploc bags accompanied by a wet paper towel to maintain humidity and minimize edge effects from evaporation. In practice, this likely contributed a small amount of oxygen (from the wet paper towel), as EcN did typically form a biofilm ring at the air-liquid interface under this condition, which was absent for larger-volume biofilm cultures in culture tubes that were incubated strictly anaerobically without humidification. Microaerobic growth was achieved by incubation inside a Heracell VIOS 160i incubator (Thermo Scientific) set to 1% O_2_ and 5% CO_2_. Aerobic biofilm experiments were incubated statically inside a standard tissue-culture incubator set to 5% CO_2_.

### Crystal violet biofilm assay

Biofilm assays were typically performed in 96-well non-tissue-culture-treated polystyrene plates (Corning, catalog no. 3370), which were incubated anaerobically as described above. Prior to each experiment, a pre-culture was performed by inoculating a single colony into 2-3 mL YCFA in a 14 mL polypropylene culture tube (Corning, catalog no. 352063) and incubating anaerobically overnight at 37°C. The pre-culture was then homogenized by pipetting and inoculated to an initial OD_600_ of 0.04 in YCFA media containing 1% DMSO or the desired bile acid diluted from a 100x stock prepared in DMSO. Each cell suspension was distributed across six wells (100 µL/well) of the 96-well plate. Three media blank wells were included for each experimental treatment (e.g. different bile acids). The plates were then incubated statically under the appropriate oxygen condition for 20-24 hours.

Following the incubation, a representative well from each condition was imaged at 4x under phase contrast on an Evos M5000 digital inverted microscope. Then, two wells from each condition were thoroughly resuspended by pipetting and OD_600_ readings for those wells were taken on a Biotek Synergy 2 plate reader. To the remaining wells, 50 µL of Bouin’s fixative (Poly Scientific, catalog no. S129) was directly added to the culture media by careful pipetting down the sides of the wells. The plates were incubated with the fixative for 15-20 min at room temperature (RT). Then, the supernatants were removed by inverting the plates over a tray and shaking sharply. The plates were washed twice by submersion in tap water followed by inversion and shaking, and were tapped against a paper towel after the second wash to remove excess liquid. Subsequently, 125 µL of 0.1% crystal violet (1:10 dilution in water of 1% crystal violet aqueous solution, Sigma-Aldrich, catalog no. V5265) was added to each well by careful pipetting down the sides of the wells. After 10 min at RT, the crystal violet solution was removed by inversion and shaking, and the plate was washed twice by successive submersion in two different trays of tap water. After removing excess liquid by tapping against a paper towel, plates were placed on top of or propped against culture tube racks and allowed to air dry upside down overnight. Once dry, the plates were imaged from the bottom using an Epson Perfection V600 scanner. Then, 125 µL of glacial acetic acid was added to each well. After 10 min at RT, the plates were gently agitated by hand to homogenize the resolubilized crystal violet and OD_550_ readings were taken on a Biotek Synergy 2 plate reader.

Larger-volume biofilm assays to illustrate the effects of oxygen or *fliC* deletion were performed in 5 mL polystyrene culture tubes (Corning, catalog no. 352058). All steps were the same as for the 96-well plate assay except that the culture volume was 1 mL, the volumes of the fixative, staining solution, and acetic acid were scaled proportionally (0.5 mL fixative, 1.25 mL of crystal violet solution, and 1.25 mL of glacial acetic acid per tube), and the washes were performed by gentle pipetting (2 mL water per tube per wash) instead of by submersion.

### Fluorescent staining and confocal microscopy of biofilms

For confocal fluorescence microscopy, biofilms were grown on 22 x 22 mm #1.5 coverslips (Corning, catalog no. 2850-22) placed in 6-well non-tissue-culture-treated polystyrene plates (VWR, catalog no. 10861-554). Inoculation from a pre-culture was performed as described above into YCFA + 100 µM LCA, and 2 mL of the resulting cell suspension was added to each well. The cultures were incubated anaerobically for 24 hours. Then, the supernatant was removed from the wells by pipetting and 2 mL of the appropriate staining solution was added by gentle pipetting down the sides of the wells. For staining eDNA, we used 2 µM TOTO-1 (Invitrogen, catalog no. T3600) in combination with 10 µM SYTO 60 (Invitrogen, catalog no. S11342) as a counterstain for intact cells. For staining proteins, we used FilmTracer SYPRO Ruby (Invitrogen, catalog no. F10318). For staining cellulose, we used Calcofluor White (Sigma-Aldrich, catalog no. 18909) at a final concentration of 0.025% (1:4 dilution) in combination with 5 µg/mL FM 1-43FX (Invitrogen, catalog no. F35355) as a marker for cells. All stains were diluted in phosphate buffered saline (PBS, with calcium and magnesium), pH 7.4, except for FilmTracer SYPRO Ruby, which is supplied at 1x concentration. Biofilms were incubated with the stains for 30 min in the dark at RT, then gently washed twice for 5 min each with 2 mL PBS by careful pipetting down the sides of the wells. After removing the second wash, the coverslip was lifted using forceps and placed biofilm side up onto a glass microscope slide. A drop of VECTASHIELD Antifade Mounting Medium (Vector Laboratories) was added on top of the biofilm and then topped with a second coverslip. The cover slips were sealed and affixed to the slide with clear nail polish. The biofilms were then immediately imaged on a PerkinElmer UltraVIEW spinning disk confocal microscope with a 40x objective. For each biofilm, a Z-stack spanning the thickness of the biofilm was collected with a step size of 0.5 µm. Image processing (consisting of taking a maximum intensity projection and adjusting the contrast and brightness) was performed in Fiji. For the SYPRO Ruby stain, we used the histogram of an unstained control biofilm imaged under identical conditions as a reference to determine the level of background signal.

For staining of flagellin, an LCA-induced biofilm was grown on a coverslip as described above. After 24 hours, the biofilm was fixed by adding 0.5 volume of Bouin’s fixative to the media. After 20 min at RT, the fixed biofilm was washed three times for 5 min each with 2 mL of PBS, then blocked with 5% normal donkey serum (Jackson ImmunoResearch Laboratories, catalog no. 017-000-121) in PBS for 6 hours at RT. The biofilm was then incubated with the anti-flagellin primary antibody (Abcam, catalog no. ab93713, 1:100 dilution in 1.5 mL blocking solution) at 4°C. After 12 hours, the biofilm was washed three times for 5 min each with 2 mL blocking solution, then incubated with donkey anti-rabbit Alexa Fluor 568 secondary antibody (Abcam, catalog no. ab175470, 1:1000 dilution in PBS) for 1 hour at RT in the dark. Finally, the biofilm was washed three times for 5 min each with 2 mL of PBS + 10 µM Hoechst 33342 (Thermo Scientific, catalog no. 62249), then mounted on a slide and imaged in the manner described above.

### Enzymatic treatment of biofilm cultures

For testing the effects of DNase and proteinase K, a biofilm assay in YCFA + 50 µM LCA was set up in a 96-well plate as described above. For one set of 6 wells, the media included 10 µg/mL DNase I (Roche, catalog no. 10104159001), while for another set of 6 wells the media included 1 mg/mL proteinase K (Qiagen, catalog no. RP107B-1). Untreated control wells contained only YCFA + 50 µM LCA. The cultures were incubated anaerobically for 24 hours and then processed as described above under “Crystal violet biofilm assay.”

### Random transposon mutagenesis and high-throughput screening

To generate random transposon insertion mutants in EcN, we used EZ-Tn5 transposase (Biosearch Technologies, catalog no. TNP92110) according to the manufacturer’s instructions. Briefly, the transposon DNA was generated by PCR amplifying the kanamycin resistance cassette from pKD4 using primers with mosaic end overhangs. Following DpnI digestion and purification with the QIAquick PCR Purification Kit (Qiagen), the PCR product was diluted to 100 ng/µL in TE buffer (10 mM Tris-HCl pH 7.5, 1 mM EDTA pH 8.0). The transposome was assembled by incubating 2 µL of the PCR product with 4 µL of EZ-Tn5 transposase and 2 µL glycerol at RT for 30 min. Electrocompetent EcN (plasmid-cured) was prepared from a 20 mL log-phase culture by two successive washes and concentration in 10% glycerol (final resuspension in 100 µL volume). The cells were incubated with 1 µL of transposome for 10 min on ice prior to electroporation. Following electroporation at standard *E. coli* settings (1.8 kV, 25 µF, 200 Ω), the cells were recovered in 1 mL SOC medium for 1.5-2 hours at 37°C with shaking at 200 rpm, then plated on LB agar with 50 µg/mL kanamycin. On average, this protocol yielded around 2000 mutants per transformation; thus, the transformation was repeated until we had generated a total of ∼10,000 mutants.

To screen the mutants for LCA-induced biofilm formation, individual colonies were picked using sterile wooden round toothpicks and inoculated into 2 mL YCFA in deep-well 96-well plates (Corning, catalog no. 3960). The plates were sealed with AeraSeal films (Excel Scientific) and the cultures were grown anaerobically at 37°C overnight, then resuspended by pipetting with a multichannel 1 mL pipette. For each plate of overnight cultures, two 96-well non-tissue-culture-treated assay plates were prepared, one containing 100 µL/well YCFA + 1% DMSO, and the other containing 100 µL/well YCFA + 100 µM LCA. The assay plates were inoculated by dipping a 96-pin disposable sterile replicator (Scinomix, catalog no. SCI-5010-OS) into the overnight culture plate, then dipping and swirling it in the assay plate. The YCFA + 1% DMSO assay plate was inoculated first to avoid any carryover of LCA. Additionally, 50 µL was transferred from each well of the overnight culture plate to a third 96-well plate containing 50 µL/well 50% glycerol, which was then stored at −70°C (thus generating a frozen stock for each transposon mutant). The inoculated assay plates were incubated anaerobically at 37°C for 20-24 hours, then processed as described above under “Crystal violet biofilm assay,” except that the wells were not imaged prior to fixation and OD_600_ readings were also not taken, as for the purposes of the high-throughput primary screen, we did not include replicate wells that could be resuspended, although we did note occasional wells with a complete failure to grow and omitted those from further analysis. Mutants were typically processed in sets of 952 (ten 96-well plates, including four wells of media blanks and four wells of EcN Δ*clbQ*::*kan* in each set as a reference for the WT biofilm phenotype). Mutants with an OD_550_ that was at least three standard deviations higher or lower than the respective control wells in either condition (1% DMSO or 100 µM LCA) were selected for secondary screening. After streaking out these mutants from the frozen stock plates onto LB + 50 µg/mL kanamycin, secondary screening was performed according to our standard crystal violet biofilm assay protocol.

### RNA-seq and RT-qPCR

For the RNA-seq time course, nine replicate 5 mL cultures of EcN were grown anaerobically to mid-log phase (OD_600_ 0.4-0.6) in YCFA. The log-phase cultures were then spiked with 1% DMSO, 100 µM LCA, or 100 µM CDCA (three cultures per condition) and transferred (1 mL/well, one well per log-phase culture) to a 12-well non-tissue-culture-treated plate (VWR, catalog no. 10861-556). Four identical plates were set up in this manner and then incubated anaerobically at 37°C. After 15, 30, 90, and 180 minutes, one plate at a time was removed from the incubator and the contents of each well were collected into microcentrifuge tubes by pipetting and scraping to collect any adherent cells along with the planktonic fraction. The collected cells were pelleted at 9000 g for 2 min, flash-frozen in liquid nitrogen after removing the supernatant, and stored at −70°C. RNA was extracted from the frozen pellets using the RNeasy Mini Kit (Qiagen) with a slightly modified protocol as previously described,^91^ including the manufacturer’s recommended on-column DNase treatment step. The extracted RNA was further treated with the TURBO DNA-*free* Kit (Invitrogen) using the manufacturer’s “rigorous” protocol to remove any remaining genomic DNA. Total RNA was quantified with the Qubit RNA HS Assay Kit (Thermo Fisher Scientific) and quality was assessed using RNA ScreenTape on TapeStation 4200 (Agilent Technologies). Sequencing libraries were generated from 400 ng of total RNA using the TruSeq Stranded Total RNA Kit (Illumina, catalog no. 20020596) in combination with the Illumina Ribo-Zero Plus rRNA Depletion Kit (Illumina, catalog no. 20037135). Libraries were quantified with the Qubit dsDNA HS Assay Kit (Thermo Fisher Scientific) and the average library size was determined using D1000 ScreenTape on TapeStation 4200 (Agilent Technologies). Libraries were pooled and sequenced on a NovaSeq 6000 (Illumina) to generate 30 million single-end 50 bp reads for each sample.

Low-quality bases were removed from the RNA-seq data using Trimmomatic (LEADING:27 TRAILING:27 SLIDINGWINDOW:4:20 MINLEN:3).^92^ Reads were aligned to the EcN reference genome (GenBank accession no. CP007799.1) using Bowtie 2,^93^ then counted with featureCounts,^94^ using the options to count all multi-overlapping reads (because reads could span two co-transcribed genes in an operon) but not multi-mapping reads. Differential gene expression analysis was performed in R (version 4.3.2) using the package DESeq2.^95^ Log_2_ fold change shrinkage was applied using the lfcShrink function in DESeq2 with the “apeglm” method. Genes were considered differentially expressed if the adjusted *p*-value was <0.05 and absolute value of the adjusted log_2_ fold change was >1. Read coverage for selected genes of interest was inspected using Integrative Genomics Viewer (IGV).^96^ For Supplementary Fig. 3F, read counts were obtained for each base in the region around *yhhT* using the “depth” command in SAMtools (version 1.7).^97^

For RT-qPCR, cultures were set up and collected in the same manner as for the RNA-seq experiment. Following RNA extraction and TURBO DNA-*free* treatment, cDNA was synthesized using the iScript cDNA Synthesis Kit (Bio-Rad) according to the manufacturer’s instructions. Then, qPCR was performed on a QuantStudio 6 Pro Real-Time PCR System (Applied Biosystems), using iTaq Universal SYBR Green Master Mix (Bio-Rad) in 20 µL reactions with 100 ng of input cDNA. Primers for qPCR were designed using IDT’s PrimerQuest tool. We used *frr* as the housekeeping reference gene for ΔΔCq quantification of relative gene expression, based on inspection of our RNA-seq data for genes with stable expression across all experimental treatments and time points. All qPCR plates included the housekeeping reference gene and no-RT controls for each sample, as well as a no-template control for each primer pair. Melt curve analysis was performed to verify the specificity of each primer pair.

### Proteomics

#### Sample collection and cell lysis

For proteomics, the biofilm cultures were set up similarly to the RNA-seq experiment, except that a 10 mL culture volume was used in 100 mm polystyrene Petri dishes in order to generate sufficient material. Three replicate cultures were set up per treatment (1% DMSO vs. 100 µM LCA vs. 100 µM CDCA). After 180 min of anaerobic incubation at 37°C, the entire culture volume was collected into 50 mL conical tubes with pipetting and scraping to capture adherent cells, and the cells were centrifuged at 4000 g for 10 min, yielding pellets with a wet weight of ∼100 mg. The cells were washed once with 25 mL PBS (pH 7.4), flash-frozen in liquid nitrogen, and lyophilized overnight to achieve complete dryness. To capture membrane-bound as well as cytoplasmic proteins, we used an acid-based cell lysis protocol adapted from precedent work,^98^ which is briefly described here. The lyophilized samples were placed at −78°C in a dry ice/acetone bath, and 300 µL of a 1:9 toluene:triflic acid mixture was added dropwise in a ventilated hood. The temperature was gradually increased to 4°C and the tubes were gently agitated by hand with intermittent venting for 60 min. The liquefied and dark cell lysis suspension resulting from the acid treatment was quenched by slowly adding 900 µL ice-cold water:methanol:pyridine (1:1:3) to the samples at −78°C (dry ice/acetone bath). To remove the strong acid and base, the samples were diluted with 10 mL of a methanol:water (1:1) solution and concentrated to a final volume of 1 mL using an Amicon 3 kDa MWCO ultra centrifugal filter (Millipore, catalog no. UFC9003). The samples were washed twice with 1 mL of 100 mM Tris-HCl, pH 8.0, then lyophilized and re-suspended in 1 mL of buffer containing 5 M urea in 100 mM Tris-HCl, pH 8.0. The total protein content was quantified using the Pierce BCA Protein Assay Kit (Thermo Scientific) and the samples were stored at −20°C until subsequent steps.

#### LC-MS/MS sample preparation

200 µg of protein was suspended in 240 µL of buffer (100 mM bicine, 5 M urea, pH 8.0). Cysteine disulfides were reduced by adding tris (2-carboxyethyl) phosphine (TCEP) to 5 mM final concentration and incubating for 15 min. The reduced samples were alkylated with chloroacetamide (25 mM, 15 min in the dark). Samples were then diluted to a final volume of 1 mL with buffer (100 mM Tris HCl, 1 mM CaCl_2_, pH 8.0) and digested into peptides using 4 µg of trypsin (Promega) overnight at 37°C. Finally, the samples were acidified with 10 µL of formic acid (10%), centrifuged (12,000 g for 5 min), and the topmost 975 µL of samples were collected.

#### TMT labeling

To prepare for TMT labeling, 300 µL of the acidified peptides were cleaned and desalted using a C18 Sep-Pak cartridge with 50 mg sorbent (Waters). The column was washed twice with 1 mL of 95% acetonitrile and 0.1% trifluoroacetic acid (TFA) and then equilibrated three times with 1 mL of 0.1% TFA. Samples were loaded into the column and washed six times with 1 mL of 0.1% TFA. Finally, peptides were eluted from the column using 0.6 mL buffer (60% acetonitrile/0.1% TFA). The eluted samples were dried using a speed vacuum and re-suspended in 100 µL of 200 mM HEPES buffer (pH 8.5). Peptides were quantified using the Pierce Quantitative Colorimetric Peptide Assay (Thermo Scientific), and 60 µg of peptides from each sample was collected and adjusted into 0.6 µg/µL concentration for further processing.

The 10-plex TMT reagents (Thermo Scientific) were equilibrated to RT. Each 0.8 mg TMT reagent vial was dissolved in 41 µL of anhydrous acetonitrile. Then, 41 µL of the TMT reagent was mixed with 60 µg of peptide samples, and labeling was allowed to occur for 1 hour at RT. Nine of the ten TMT tags (TMT-126 to TMT-130C) were used for the nine samples. A 2 µL aliquot from each sample was taken and mixed for a test run, while the remaining samples were stored at −80 °C until the test samples were processed. The mixed test samples were dried and re-suspend to a final volume of 20 µL of 0.1%TFA buffer. They were then cleaned using GL-Tip stage tips (GL Sciences, catalog no. 7820-11200), and analyzed on an Orbitrap Fusion Lumos Tribrid mass spectrometer (Thermo Fisher Scientific). This analysis was performed to evaluate the quality of the TMT labeling and to normalize the labeling efficiency. Upon successful completion of the test run, the main samples stored at −80 °C were subjected to a freeze-thaw cycle. The TMT labeling reaction was quenched by adding 8 µL of 5% hydroxylamine to each sample, followed by 15 min incubation. Based on the test run result, the samples were normalized, combined, and dried by speed vacuum for the subsequent steps.

#### Peptide fractionation

The dried mixed samples were resuspended in 1 mL of 0.1% TFA and then purified using a C18 Sep-Pak cartridge containing 50 mg of sorbent (Waters). Initially, the C18 column was washed twice with 1 mL of a buffer (95% acetonitrile and 0.1% TFA) and then equilibrated three times with 1 mL of 0.1% TFA. The samples were subsequently loaded onto the column and washed three times with 1 mL of 0.1% TFA. Finally, peptides were eluted from the column using 0.6 mL of eluting solution (60% acetonitrile/0.1% TFA). The eluted peptides were dried using a speed vacuum and then re-suspended in 300 µL of 0.1% TFA solution.

The purified samples were fractionated using a high pH reversed-phase peptide fractionation column (Thermo Scientific, catalog no. 84868). Initially, the column was conditioned using acetonitrile and 0.1% TFA solution. The sample was then loaded onto the column and eluted using eight different elution solutions prepared with varying acetonitrile concentrations, ranging from 10% to 50%. The eight fractions of eluted peptides were collected and dried. These fractions were further cleaned up again using a C18 Sep-Pak cartridge containing 50 mg of sorbent (Waters), following the same procedure described above. The fractionated and cleaned samples were resuspended with 15 µL of 2% acetonitrile and 0.1% formic acid. Finally, the total peptides were quantified using the Pierce Quantitative Colorimetric Peptide Assay (Thermo Scientific).

#### LC-MS/MS analysis

An LC-MS/MS analysis was performed for each fraction using an Orbitrap Eclipse Tribrid mass spectrometer (Thermo Fisher Scientific), which was equipped with FAIMS (Field Asymmetric Ion Mobility Spectrometry). The samples were analyzed at 40 and 60 CV. The mass spectrometer was paired with a Dionex Ultimate 3000 RSLCnano Proflow system (Thermo Fisher Scientific) that contained an Aurora Series 25 cm x 75 µm I.D. column (IonOpticks). A low pH reversed-phase separation was conducted using a mass spectrometry grade solvent A (2% acetonitrile, 0.1% formic acid in 98% water) and solvent B (0.1% formic acid, 98% acetonitrile, 2% water) over a total duration of 185 min. The process began by ramping solvent B from 2% to 4% at a flow rate of 400 nL/min for 9.9 minutes. The gradient was held at 4% for 0.1 min, during which the flow rate was reduced from 400 nL/min to 300 nL/min. For the next 135 min, the gradient of solvent B was increased from 4% to 30% at a flow rate of 300 nL/min. Maintaining the same flow rate, the gradient was then ramped from 30% to 75% over 15 min, and subsequently increased from 75% to 90% during the following 3 min. Solvent B was held at 90% for 6.9 min. The flow rate was then switched from 300 nL/min back to 400 nL/min, and the gradient was reduced from 90% to 2% for 0.1 minutes. Finally, the system was maintained at 2% solvent B at a flow rate of 400 nL/min for 15 min, completing the run.

For these analyses, a FASTA file of the *E. coli* Nissle 1917 database UP000263938 was created from UniProt (downloaded in November 2023). The mass spectrometry run was searched in real-time against this database using an in-house instrument API program called inSeqAPI.^99^ Protein-closeout was employed stopping analysis of peptides from proteins with 3 distinct peptides, 5 unique peptides, or 10 peptide spectral matches already quantified. The analysis of intact peptides began with the MS1 scan, which utilized an Orbitrap mass detector set to a resolution of 120,000 and a normalized AGC target of 250%. The scan range was defined between m/z 350 and 1350, ensuring a comprehensive analysis of the mass-to-charge ratios of the peptides. During this process, the system used a 30% RF lens to efficiently focus the ions. The instrument operated in data-dependent mode, where it selected up to 10 of the most intense precursor ions that were subjected to MS2 fragmentation. The maximum injection time for the MS1 analysis was set to 50 ms, allowing sufficient time for ion accumulation before the MS2 fragmentation stage. After the selection of peptides for MS2 fragmentation, the chosen precursor ions were isolated using a quadrupole and fragmented using collision-induced dissociation (CID) with a collision energy set at 35% and a CID activation time of 10 ms. The resulting fragment ions were subsequently analyzed using an ion trap. During this MS2 analysis, a normalized AGC target was set at 150%, and the maximum injection time was 100 ms. Synchronous precursor selection (SPS) MS3 scans were conducted using the Orbitrap mass detector at a resolution of 50,000. The eight most intense ions from the MS2 spectrum were selected for HCD fragmentation, utilizing a normalized collision energy of 45%. This process was optimized with an AGC target of (3.0 x 10^5^) and a maximum injection time of 400 ms.

#### Proteomics data analysis

The raw files were searched using COMET (Advanced) 2019.01^100^ against a concatenated target-decoy database of the *E. coli* Nissle 1917 proteins containing common contaminants. The COMET search parameters included complete trypsin cleavage with an allowance for one missed cleavage event. A peptide mass tolerance of 20 ppm was used. For low resolution MS/MS of the fragment ion, bin tolerance and bin offset for the COMET search were tuned to 1.0005 and 0.4, respectively, to ensure optimized peptide identification. The static modifications of the peptides included carbamidomethylation of cysteine, which adds a mass of +57.0215 Da. Additionally, the static modification of the TMT tags were applied to lysine residues and the peptide N-terminus, adding a mass of +229.1629 Da. Moreover, the COMET search permitted variable modification of methionine oxidation (+15.9949 Da) and included the modification of the TMT-labeled tyrosine (+229.1629 Da). The searched peptides were filtered based on peptide-spectrum matches (PSMs) using a false discovery rate (FDR) of 1%. TMT reporter ions generated by the TMT tags were quantified using an in-house software package called Mojave.^101^ This was done by identifying the highest peak within 20 ppm of the theoretical reporter mass windows and correcting for isotope purities.

The downstream analysis was performed using R (version 4.1.0) with the MSstatTMT package (version 2.0.1).^102^ Peptide-spectrum match (PSM)-level quantifications were aggregated to the protein level, where differential abundance analysis was conducted. In the group comparisons of each treatments, the MSstatTMT package quantified the log_2_ fold change and estimated the standard error for each protein across treatment groups. In order to test the two-sided null hypothesis of no changes in abundance, model-based test statistics were compared against the Student’s t-distribution, with degrees of freedom tailored to each protein and dataset. Proteins were considered differentially expressed if the adjusted *p*-value was <0.05 and absolute value of the log_2_ fold change was >1.

### Soft-agar swimming assay

To test the swimming ability of EcN, soft agar plates were prepared by combining equal volumes of 0.6% molten agar and YCFA prepared at 2x concentration (for a final concentration of 0.3% agar, 1x YCFA) and pipetting 5 mL into each well of a 6-well plate. 100 µM LCA or 1% DMSO was included in the media when testing the impact of LCA on swimming behavior, but not when testing the swimming ability of mutants. The WT or mutant strains were inoculated individually into separate wells (three replicates per strain or condition) by dipping a sterile wooden round toothpick into 100 µL of overnight YCFA culture in a microcentrifuge tube, then stabbing the toothpick 1-2 mm into the center of the soft agar, without touching the bottom of the well. The plates were incubated aerobically, lid side up, in humidified plastic boxes at 37°C for 8 hours, after which the diameter of the cloud of swimming cells was measured with a ruler.

### Scanning electron microscopy

The samples for SEM were obtained by culturing EcN overnight in 15 mL conical tubes with 5 mL anaerobic YCFA containing 1% DMSO or 50 µM LCA. The bacterial aggregates naturally settled to the bottom of the tube and the supernatants were carefully removing without disturbing the settled aggregates, then the cells were fixed in 1 mL Karnovsky fixative (2.5% glutaraldehyde, 2% formaldehyde, 0.1 M sodium cacodylate buffer, pH 7.4) and stored at 4°C until further processing. The fixed bacterial samples were adsorbed for 5 min, at RT, to poly-lysine coated glass slides, rinsed in water, and then stained and postfixed in 1% aqueous osmium tetroxide for 15 min at RT. The slides were then rinsed in water and stained with 2% uranyl acetate for 15 min at RT. The samples were then rinsed in water and dehydrated in an ascending series of ethanol and finally incubated two times in 100% hexamethyldisilizane for 5 min each. The samples were then air dried and sputter coated with palladium / gold (5 nm thickness). The samples were imaged with a Gemini300 scanning electron microscope (Zeiss) at 1 keV using the “in-lens” secondary electron detector. Images were acquired at magnifications from 500x to 25,000x.

### Antibiotic tolerance assay

Prior to testing the effect of LCA and biofilm formation on the antibiotic tolerance of EcN, the MIC of ciprofloxacin was established by performing a microtiter broth dilution assay according to a standard clinical protocol,^103^ except that YCFA + 1% DMSO and YCFA + 100 µM LCA were used as the media instead of Mueller-Hinton broth. In both conditions, the MIC was 0.0156 µg/mL. For the antibiotic tolerance assay, a 96-well non-tissue-culture-treated plate was set up with twelve replicate wells per condition (YCFA + 1% DMSO vs. YCFA + 100 µM LCA, 100 µL/well), inoculated to OD_600_ 0.04 from an overnight YCFA culture. Unused wells were filled with water and the plate was incubated anaerobically at 37°C in a Ziploc bag humidified with a wet paper towel. Meanwhile, the remaining overnight culture was centrifuged at 20,800 g for 2 min, and the supernatant was passed through a 0.22 µm PES syringe filter, generating spent media for subsequent steps. The spent media was stored overnight in the anaerobic chamber. The following day, after 24 hours of incubation, the 96-well plate and the spent media were brought out of the anaerobic chamber. 90 µL of supernatant was transferred from each well to a new 96-well plate, representing the planktonic fraction. From the original wells, which represented the biofilm fraction, any remaining supernatant was carefully removed and discarded without disturbing any biofilm present. To a second new 96-well plate, 90 µL/well of spent media + 1% DMSO or 90 µL/well of spent media + 100 µM LCA was added (three wells per condition). Then, 10 µl of 5 µg/mL ciprofloxacin (0.5 µg/mL final concentration) was added to either three wells or six wells per condition, respectively, in the spent media and planktonic fraction 96-well plates, pipetting well to mix. The contents of the spent media 96-well plate were then gently transferred to the respective wells in the original 96-well plate, thus treating the biofilm fraction with the antibiotic. The use of spent media as the carrier for the biofilm antibiotic treatment was to ensure that both the biofilm and the planktonic fractions would remain in stationary phase during the antibiotic treatment. Additionally, three wells per condition of the antibiotic-treated planktonic fraction were transferred back to the respective wells in the original plate, representing combined treatment of both the planktonic and biofilm fractions. Finally, three wells per condition of the untreated (no antibiotic added) planktonic fraction were transferred back to the respective wells in the original plate, thereby recombining the untreated planktonic and biofilm cells into a single population. These wells along with the remaining three untreated wells per condition of separated planktonic cells and biofilm cells were thoroughly resuspended by pipetting (resuspension was in PBS for the biofilm fraction) and plated for CFUs on LB agar to represent the starting number of CFUs in each condition (the reference point for 100% survival).

Both of the antibiotic-treated 96-well plates (one containing the biofilm fraction and combined biofilm + planktonic fractions, the other containing the separated planktonic fraction) were incubated anaerobically at 37°C for another 24 hours. Then, all wells were thoroughly resuspended and plated for CFUs on LB agar to assess the number of surviving cells. CFUs were counted after 24 hours of incubation at RT, and the surviving fraction was reported as the number of CFUs post-antibiotic treatment divided by the number of CFUs obtained from the untreated controls for each condition at the start of the antibiotic treatment.

### Strain competition assay

For the strain competition experiment, overnight anaerobic YCFA cultures of each strain were diluted to ∼5 x 10^6^ CFU/mL in YCFA + 1% DMSO or YCFA + 100 µM LCA. The competing strains were set up in pairwise combinations with EcN Δ*clbQ*::*kan* with an equal inoculum of each strain. For each combination and treatment, 1 mL/well was dispensed across three replicate wells of a 12-well non-tissue-culture-treated plate (VWR, catalog no. 10861-556). The cultures were incubated anaerobically at 37°C. Meanwhile, growth curves were set up for each strain individually by diluting to the same inoculum in both YCFA + 1% DMSO and YCFA + 100 µM LCA, with six replicate wells per condition (100 µL/well) in a 96-well non-tissue-culture treated plate. The growth curve plate was incubated at 37°C in a Biotek Epoch 2 plate reader inside an anaerobic chamber, with continuous double-orbital shaking at 282 cpm. OD_600_ readings were taken every 15 min for 24 hours, utilizing the pathlength correction function of the plate reader.

After 24 hours of incubation of the co-cultures, the supernatants (representing the planktonic fraction) were transferred to fresh 12-well plates, homogenized by vigorous pipetting, and plated for CFUs on both LB agar (total CFU count for both strains) and LB agar + 50 µg/mL kanamycin (CFU count for EcN). The original wells (representing the biofilm fraction) were washed gently once with 1 mL PBS to remove residual planktonic cells, then resuspended in 1 mL PBS with vigorous pipetting and scraping to thoroughly disrupt any biofilm, and plated for CFUs on LB agar and LB agar + 50 µg/mL kanamycin. CFUs were counted after 24 hours of incubation at RT. The proportion of EcN was calculated as the number of CFUs counted on LB agar + 50 µg/mL kanamycin divided by the number of CFUs counted on LB agar.

### Human intestinal organoid cultures

Human intestinal organoids were derived as previously described from colonic tissue procured by Donor Network West from a deidentified post-mortem donor.^104^ After establishment of the organoid line, organoids were routinely grown in 3D culture according to established protocols^105^ in Cultrex Reduced Growth Factor Basement Membrane Extract, Type 2 (R&D Systems, catalog no. 3533-005-02) or Matrigel Growth Factor Reduced Basement Membrane Matrix (Corning, catalog no. 356231), dispensed as 14-15 µL droplets for a total of 100 µL/well in 12-well non-tissue-culture-treated plates or 200 µL/well in 6-well non-tissue-culture-treated plates. The droplets were overlaid with expansion medium (1 mL/well for 12-well plates, 2 mL/well for 6-well plates) consisting of Advanced DMEM/F12 (Gibco, catalog no. 12634028), 10 mM HEPES buffer (pH 7.0; Gibco, cat. no. 15630-056), 1x GlutaMAX supplement (Gibco, cat. no. 35050-038), 1x B-27 supplement (Gibco, catalog no. 17504044), 1.25 mM N-acetylcysteine (Sigma-Aldrich, catalog no. A9165), 10 mM nicotinamide (Sigma-Aldrich, catalog no. N0636), 100 ng/mL recombinant human Noggin (R&D Systems, catalog no. 6057-NG), 50 ng/mL recombinant human epidermal growth factor (R&D Systems, catalog no. 236-EG), 5 nM gastrin (Tocris, catalog no. 3006), 0.5 µM ALK5 inhibitor A83-01 (Tocris, catalog no. 2939), 10 µM p38 inhibitor SB202190 (Sigma-Aldrich, catalog no. S7076), 250 ng/mL recombinant human R-Spondin 3 (RSPO3; produced in-house or obtained from R&D Systems, catalog no. 3500-RS), 1 µM prostaglandin E2 (PGE2; Tocris, catalog no. 2296), 0.5 nM Wnt surrogate-Fc fusion protein (Gibco, catalog no. PHG0402), and 5% heat-inactivated fetal bovine serum (FBS, Gibco A5209501). This expansion medium was generally prepared at 2x concentration with all ingredients except RSPO3, PGE2, Wnt surrogate, and FBS, and was stored at −20°C for up to four months. RSPO3, PGE2, Wnt surrogate, and FBS were added after thawing the 2x media and diluting to 1x concentration with Advanced DMEM/F12, after which the complete medium was stored at 4°C and used within one month.

Organoids were passaged every 5-7 days by harvesting in 12 mL of ice-cold Advanced DMEM/F12++ (Advanced DMEM/F12 containing 10 mM HEPES and 1x GlutaMAX) per 600 µL of Cultrex or Matrigel, centrifuging at 400 g for 5 min at 4°C, resuspending in 1 mL ice-cold Advanced DMEM/F12++, and triturating 15 times with a 1 mL pipette, followed by 30 times with a P10 tip attached to the end of the P1000 tip. Alternatively, following the first centrifugation step, organoids were resuspended on ice in 1 mL of TrypLE Express (Gibco, catalog no. 1260413) containing 10 µM ROCK inhibitor Y-27632 (STEMCELL Technologies, catalog no. 72304), then incubated in a 37°C water bath for 1-2 minutes, and triturated 10 times with a P1000 pipette followed by a P10 tip attached to the P1000 tip. Organoids were passaged typically at a 1:3 or 1:4 ratio, such that every ∼1 mm of pelleted organoid fragments was resuspended in 600 µL of Cultrex or Matrigel and seeded across half of a multi-well plate. After seeding, the plates were incubated upside down for 30 min at 37°C to allow the droplets to solidify, then the droplets were overlaid with expansion medium containing 10 µM Y-27632. After 2-3 days, the expansion medium was replenished without Y-27632, and the medium was refreshed every 2-3 days thereafter until the next passage. Organoids were used for experiments between passages 10-20.

### Bacterial co-cultures with human colonic organoid-derived monolayers

To facilitate access to the apical side of the intestinal epithelium for bacterial co-cultures, organoids were cultured as 2D monolayers as follows, using a protocol adapted from Gonyar et al.^55^ 5-7 days after organoid passaging, transparent polyester membrane 24-well (6.5 mm) cell culture inserts with a 0.4 µm pore size (Corning, catalog no. 3470) were coated with 50 µL/insert of a 1:30 dilution in water of 1 mg/mL human collagen IV (Sigma-Aldrich, catalog no. C5533) stock solution prepared in 100 mM acetic acid. The inserts were incubated with the collagen solution for 1 hour at RT followed by 1 hour at 37°C. Meanwhile, organoids were collected according to the above protocol, except that after the initial centrifugation step, the organoids were resuspended in 3 mL TrypLE Express containing 10 µM Y-27632 and incubated in a 37°C water bath for 10 min, followed by trituration 10 times with a P1000 pipette followed by a P10 tip attached to the P1000 tip. The dissociation progress was checked under a phase contrast microscope at 4x magnification and the 37°C incubation was repeated in 5 min intervals, followed by trituration, until the organoids had dissociated into mostly single cells with a few remaining small clusters of cells (typically a total of 25-30 min at 37°C).

Following the dissociation, TrypLE Express was inactivated by adding 9 mL of ice-cold Advanced DMEM/F12++ containing 10 µM Y-27632 and the cells were collected by centrifugation at 400 g for 5 min at 4°C. The pellet was resuspended in 1 mL of Advanced DMEM/F12++ containing 10 µM Y-27632 and the cells were counted in a hemocytometer using Trypan Blue (Gibco, catalog no. 15250061) to assess viability (typically >90%). The centrifugation was repeated and the cells were resuspended in expansion medium containing 10 µM Y-27632 to a concentration of 10^6^ cells/mL. Then, 100 µL of the cell suspension was seeded onto each insert (100,000 cells/insert) after removing excess collagen solution, and 0.5 mL of expansion medium containing 10 µM Y-27632 was added to the basolateral chamber of each well of the 24-well plate. The cultures were incubated for two days before replenishing the expansion medium without Y-27632. The medium was replenished again on the fourth day. After five days in expansion medium, monolayers were fully confluent and were switched to differentiation medium, consisting of the same recipe as expansion medium but omitting nicotinamide, SB202190, and Wnt surrogate. The differentiation medium was replenished every two days.

On the fourth day of differentiation, an overnight culture of EcN pMut1-sfGFP was inoculated in YCFA and incubated aerobically (to enable proper folding of GFP). On the fifth day of differentiation, the basolateral media was removed and the apical surface of the monolayers was washed gently three times with 100 µL Dulbecco’s PBS (DPBS, with calcium and magnesium; Gibco, catalog no. 14040141). EcN pMut1-GFP was inoculated to OD_600_ 0.04 (∼3-4 x 10^7^ CFU/mL) in 75% YCFA, 25% Advanced DMEM/F12, supplemented with 1% DMSO or 100 µM bile acid (LCA or CDCA) as appropriate. 100 µL of the EcN suspension or media blank was added to the apical side of each monolayer (with three replicate wells per condition), then the basolateral compartment was filled with 0.5 mL of differentiation medium containing 1% DMSO or 100 µM bile acid as appropriate. Importantly, the differentiation medium used during the co-culture (but not prior to that) was prepared with a homemade version of B-27 supplement with 100-fold lower bovine serum albumin (BSA) compared to the original recipe described by the Hanna Lab, Weizmann Institute of Science (https://www.weizmann.ac.il/molgen/hanna/sites/molgen.hanna/files/users/user52/HANNA-LAB-B22-B27-PROTOCOL-V3.pdf, accessed in March 2023). The use of reduced-BSA B-27 was necessary for the co-cultures as we discovered in preliminary experiments that the high amount of BSA in typical organoid media inhibits biofilm formation by EcN. Briefly, our recipe contained the following per 50 mL of 50x supplement: Neurobasal medium (Gibco, catalog no. 21103049) to a total final volume of 50 mL; 62.5 mg BSA fraction V fatty acid free (Bio-World, catalog no. 22070023-3); 6.25 mg biotin (Sigma-Aldrich, catalog no. B4639) dissolved in 100 µL Neurobasal medium; 6.25 mg catalase (Sigma-Aldrich, catalog no. C40) dissolved in 600 µL Neurobasal medium; 625 µL of 30K units of superoxide dismutase (Sigma-Aldrich, catalog no. S5395) dissolved in 1 mL Neurobasal medium; 2.5 mg glutathione (Sigma-Aldrich, catalog no. G6013); 5 mg of L-carnitine (Sigma-Aldrich, catalog no. C0283) dissolved in 500 µL Neurobasal medium; 37.5 mg D-galactose (Sigma-Aldrich, catalog no. G0625); 40.25 mg putrescine (Sigma-Aldrich, catalog no. P5780); 2.5 µL ethanolamine (Sigma-Aldrich, catalog no. E9508); 15.75 µL of progesterone (Sigma-Aldrich, catalog no. P8783) prepared as a 1 mg/mL stock solution in ethanol; 0.5 µL of T3 (Sigma-Aldrich, catalog no. T6397) prepared as a 100 mg/mL stock solution in DMSO followed by 1:50 dilution with ethanol; 25 µL of linoleic acid (Sigma-Aldrich, catalog no. L1012) prepared as 100 mg/mL stock solution in ethanol; 25 µL of linolenic acid (Sigma-Aldrich, catalog no. L2376) prepared as 100 mg/mL stock solution in ethanol; 12.5 µL retinyl acetate (Sigma-Aldrich, catalog no. R7882) prepared as 20 mg/mL stock solution in ethanol; 25 µL D-alpha tocopherol (Sigma-Aldrich, catalog no. T3251) prepared as 100 mg/mL stock solution in ethanol; and 25 µL D-alpha tocopherol acetate (Sigma-Aldrich, catalog no. T3001) prepared as 100 mg/mL stock solution in ethanol. The ingredients were allowed to dissolve and equilibrate overnight at 4°C without agitation. Additionally, when this version of reduced-BSA homemade B-27 was used, the differentiation medium was supplemented with Insulin-Transferrin-Selenium (ITS) supplement (Gibco, catalog no. 41400045) at 1x final concentration (original B-27 supplement already contains insulin, transferrin, and selenium).

After inoculation with EcN pMut1-GFP, the co-cultures were incubated at 37°C under 1% O_2_ using a Heracell VIOS 160i incubator (Thermo Scientific). After 7 hours, Hoechst 33342 dye (Thermo Scientific, catalog no. 62249) was added to the basolateral media at a final concentration of 10 µg/mL to stain the epithelial cells, then the cultures were returned to the hypoxic incubator. At 8 hours after the start of the experiment, 50 µL of basolateral media was collected from each well and used in an LDH release assay for assessment of toxicity (Promega, catalog no. G1780). Meanwhile, the apical surface of each well was washed gently three times with 100 µL DPBS to remove non-adherent EcN, then the wells were imaged at 4x magnification under phase contrast and in the DAPI (for Hoechst 33342) and GFP channels on an Evos M5000 digital inverted microscope. Finally, the wells were lysed by adding an equal volume of 1% Triton X-100 directly to the apical chamber, incubating for 5 min at RT, and resuspending by vigorous pipetting. The lysates were transferred (150 µL/well) to a black-walled, clear-bottomed 96-well plate (Corning, catalog no. 3603) and GFP fluorescence was recorded on a Biotek Synergy 2 plate reader.

### Staining of colonic organoid-derived monolayers for MUC2

Colonic organoid-derived monolayers were seeded and differentiated as described above. On day 5 of differentiation, the monolayers were fixed in 4% paraformaldehyde for 20 min at RT, then washed twice with PBS. The fixed monolayers were permeabilized and blocked with 100 µL/well 0.5% Triton X-100, 5% normal donkey serum in PBS in the apical chamber for 1 hour at RT, then incubated with anti-MUC2 antibody clone CCP58 (1:100 dilution; Sigma-Aldrich, catalog no. MABF1989) in blocking buffer overnight at 4°C. This antibody primarily targets the intracellular form of MUC2. The following day, the monolayers were washed three times with PBS in both the apical and basolateral chambers, then stained with 100 µL/well donkey anti-mouse Alexa Fluor 568 secondary antibody (1:500; Life Technologies, catalog no. A10037) in blocking buffer overnight at 4°C. Finally, the monolayers were washed three times with PBS before imaging on an Evos M5000 digital inverted microscope.

## DATA AVAILABILITY

The raw data from the RNA-seq experiment have been submitted to the NCBI Sequence Read Archive under the accession number GSE297446, while the raw files for each peptide fraction from the proteomic experiment have been deposited into the MassIVE repository with the identifier MSV000097561.

## ACKNOWLEDGEMENTS

We thank Natalie Omattage-Huynh for generating the plasmid-cured EcN, EcN Δ*clbQ*::*kan*, and EcN pMut1-sfGFP strains; Loryn Holokai for deriving the colon organoids from primary tissue; Dennis Wolan for advice on proteomics; the next-generation sequencing core facility at Genentech for RNA-seq library preparation and sequencing; and all members of the Department of Infectious Diseases and Host-Microbe Interactions at Genentech for helpful discussions. Additionally, we are deeply grateful for the cooperation of Donor Network West and to the organ donor and their family for their generous donation enabling the advancement of scientific knowledge. This study received funding from Genentech, Inc. The funder was not involved in the study design, collection, analysis, interpretation of data, the writing of this article or the decision to submit it for publication.

## AUTHOR CONTRIBUTIONS

EP and M-WT conceived the study. BU assisted with organoid experiments and edited the manuscript. The proteomics methods and analysis were designed by CR and executed by EZ. MR performed the electron microscopy sample preparation and imaging. EP designed and performed all other experiments, analyzed the data, and wrote the manuscript. All authors read and approved the final manuscript.

