## Supplementary Information for "A microbiota-derived bile acid modulates biofilm formation by the probiotic strain *Escherichia coli* Nissle 1917"

**Supplementary Table 1: Bile acid concentrations in the water fraction of human cecal and fecal contents**

| Bile acid | Concentration (µM) | Sample type | Reference |
|---|---|---|---|
| TCDCA | Median <1 | Cecal contents | Hamilton et al., 2007 |
| GCDCA | Median <1 | Cecal contents | Hamilton et al., 2007 |
| CDCA | 0-19 | Fecal water | Van Faasen et al., 1993 |
| UDCA | 2-48 | Fecal water | Van Faasen et al., 1993 |
| LCA | 6-97 | Fecal water | Van Faasen et al., 1993 |
| TCA | Median <1 | Cecal contents | Hamilton et al., 2007 |
| GCA | Median <1 | Cecal contents | Hamilton et al., 2007 |
| CA | 0-83 | Fecal water | Van Faasen et al., 1993 |
| DCA | 4-247 | Fecal water | Van Faasen et al., 1993 |

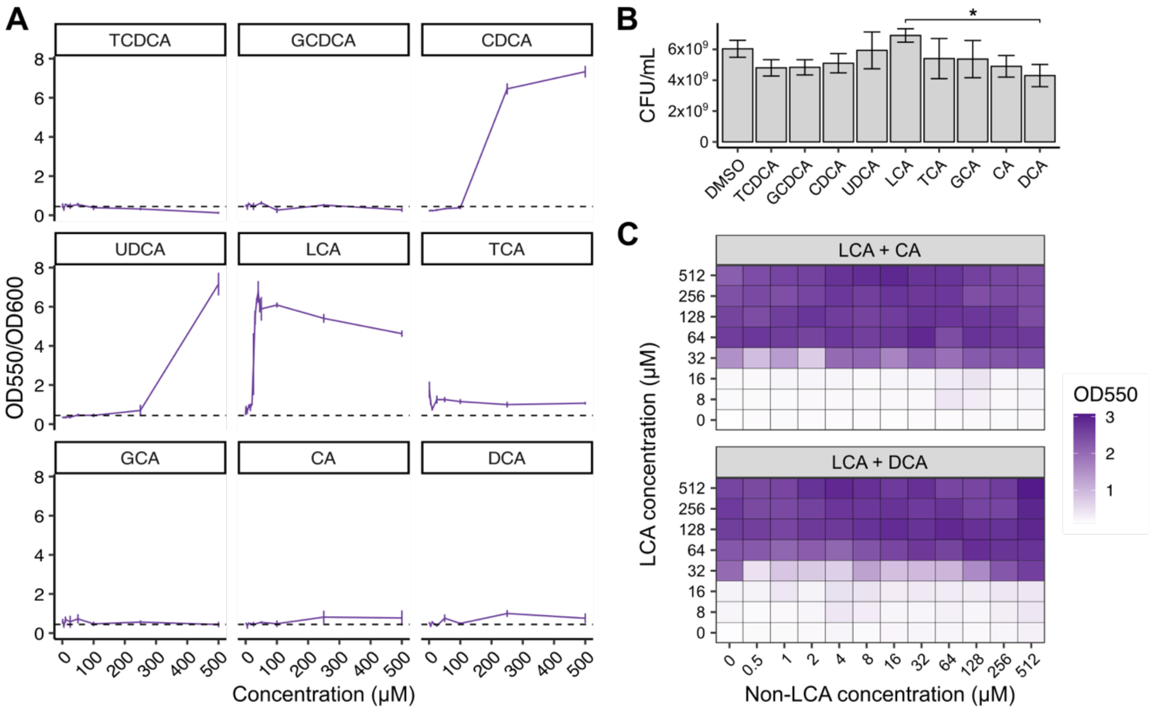


**Supplementary Fig. 1:** (A) Quantification of biofilm formation (crystal violet absorption at $OD_{550}$ normalized to $OD_{600}$ as a proxy for growth) after 24 hrs of treatment with individual bile acids at 1, 5, 10, 25, 50, 100, 250, and 500 µM. Error bars mark the standard deviation ($n = 4$ replicate wells from a representative experiment). (B) CFU counts for EcN grown in the presence of different bile acids at 500 µM. * $p < 0.05$ (ANOVA followed by Tukey HSD). None of the bile acid conditions were significantly different from the DMSO solvent control. (C) Effect of LCA paired with CA or DCA at concentrations up to 512 µM each, as tested in the style of a checkerboard assay. Each shaded box represents a unique pairwise combination of concentrations and the intensity of the color represents the measured OD550 (crystal violet absorption) for that well. Results shown are representative of two independent experiments.

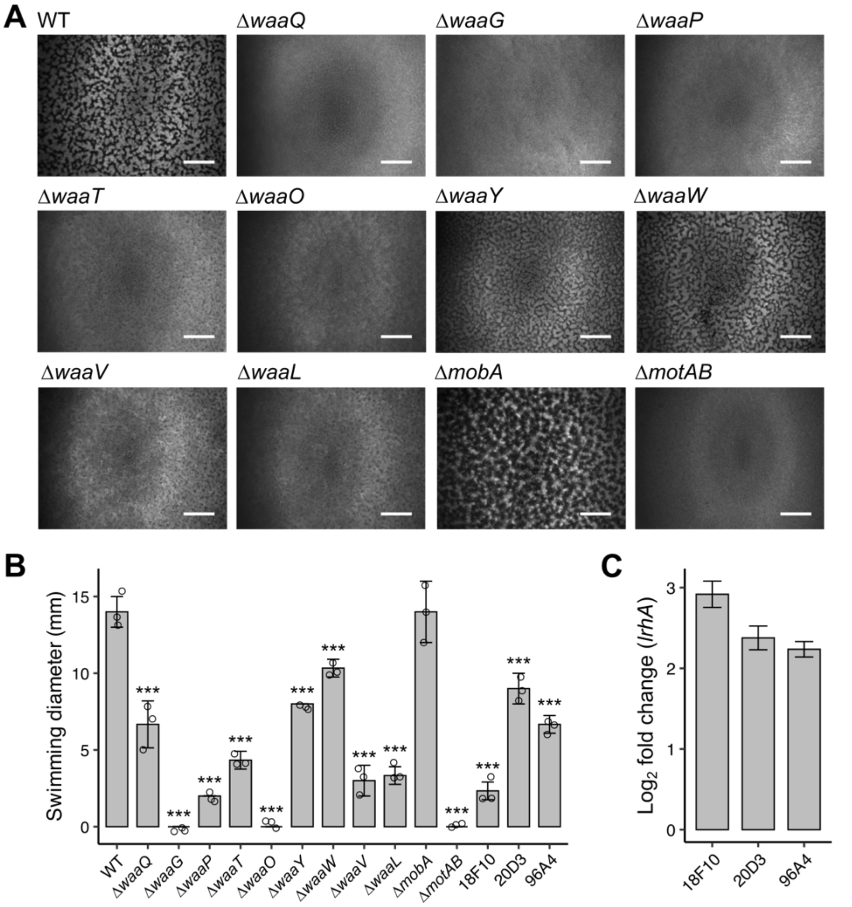


**Supplementary Fig. 2:** (A) Phase contrast images of cultures grown in YCFA + 1% DMSO in a 96-well plate for 24 hrs. Scale bars = 500 µm. (B) Diameter of the halos formed by different mutants vs. the WT after 8 hrs in a soft-agar swimming assay. *** $p < 0.001$ (Dunnett's test, comparing each mutant strain to the WT; $n = 3$). (C) RT-qPCR assessment of the expression of *lrhA* in the three transposon mutants (named by the plate and well number from which they were isolated during the screen) with Tn5 insertions in the non-coding region upstream of *lrhA*. $Log_2$ fold change was calculated relative to the WT.

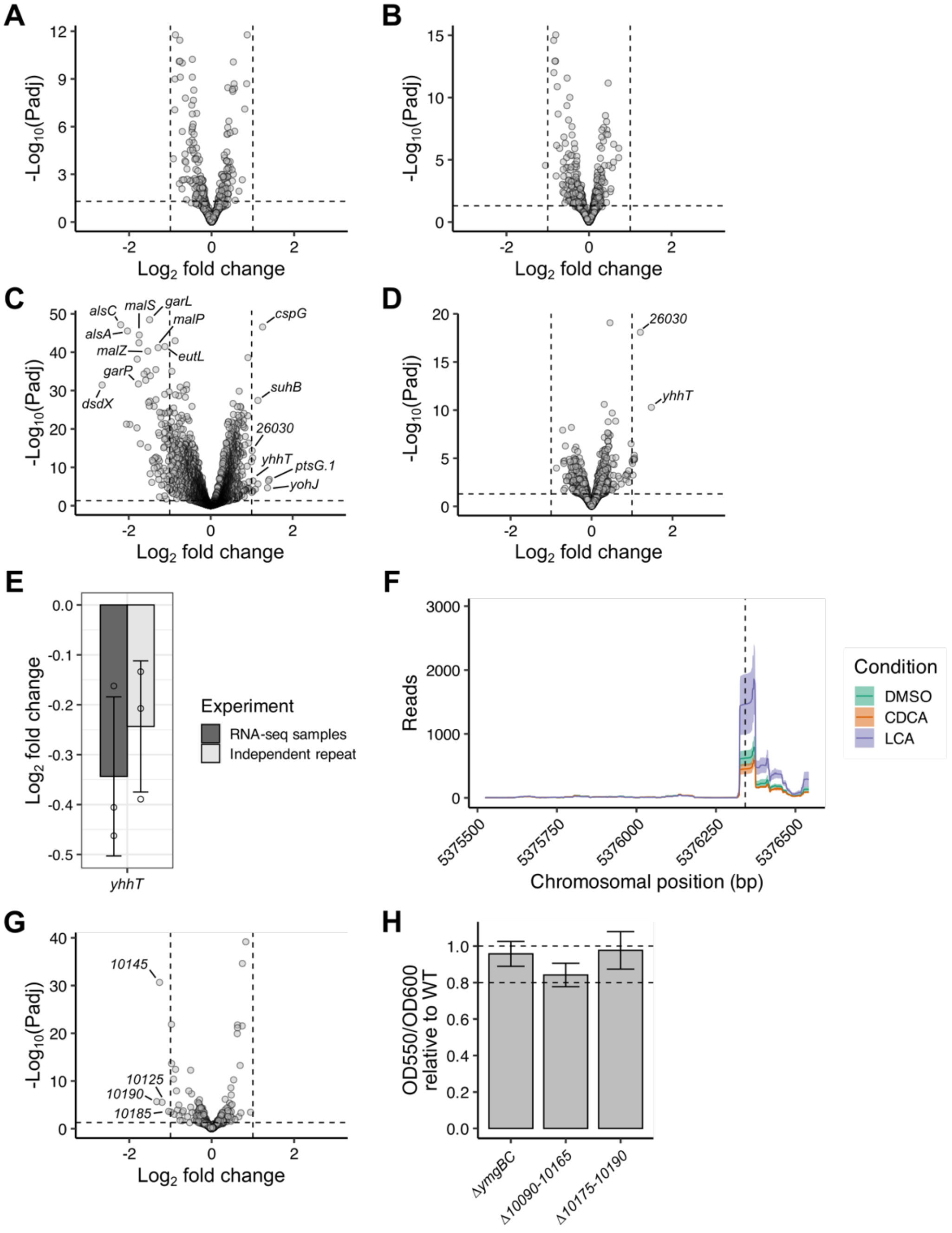


**Supplementary Fig. 3:** (A) Volcano plot for the LCA vs. DMSO RNA-seq for samples collected at 15 min. The dashed vertical lines mark a $\log_2$ fold-change cutoff of +/- 1, while the dashed horizontal line marks an adjusted *p*-value cutoff of 0.05. (B) Volcano plot for the LCA vs. DMSO RNA-seq for samples collected at 30 min. The dashed vertical lines mark a $\log_2$ fold-change cutoff of +/- 1, while the dashed horizontal line marks an adjusted *p*-value cutoff of 0.05. (C) Volcano plot for the LCA vs. DMSO RNA-seq for samples collected at 90 min. The dashed vertical lines mark a $\log_2$ fold-change cutoff of +/- 1, while the dashed horizontal line marks an adjusted *p*-value cutoff of 0.05. (D) Volcano plot for the LCA vs. CDCA RNA-seq for samples collected at 90 min. The dashed vertical lines mark a $\log_2$ fold-change cutoff of +/- 1, while the dashed horizontal line marks an adjusted *p*-value cutoff of 0.05. (E) RT-qPCR assessment of the expression of *yhhT* in EcN treated with LCA vs. DMSO for 90 min. RT-qPCR was performed both with a fresh set of samples collected independently from the RNA-seq experiment, as well as with the original RNA-seq samples, to confirm the discrepancy between RT-qPCR and the RNA-seq results. Fold changes were calculated with respect to the DMSO-treated control. Error bars represent the standard deviation (*n* = 3 replicates). (F) RNA-seq read counts along the length of *yhhT* (chromosomal positions 5375524-5376342) and 200 bp immediately downstream of the gene. The dashed vertical line represents the end of the *yhhT* coding sequence. Solid lines represent the mean read count in each condition while the shaded area represents the standard deviation.

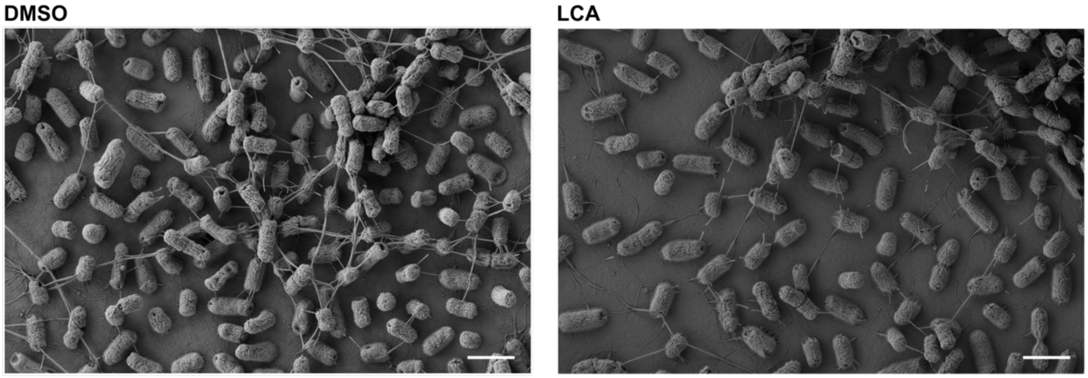


**Supplementary Fig. 4:** Representative SEM images of EcN cultures grown in YCFA + 1% DMSO or YCFA + 100 µM LCA for 24 hrs. Scale bars = 2 µm.

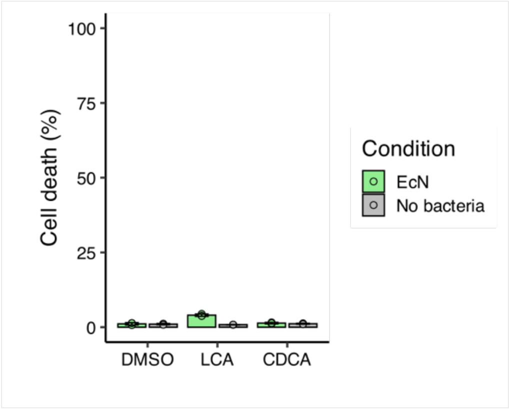


**Supplementary Fig. 5:** Epithelial cell death during co-cultures with EcN and bile acids, as measured by an LDH release assay. The signal was normalized to a “100% death” control generated by treating a well with lysis buffer provided by the assay manufacturer.

**Supplementary Table 2: Primers used in this study.**

| Primer | Sequence |
|---|---|
| pMut1sfGFP fwd | CGTTTTTATTGGTGAGAATGCGCTGAACGCGATTCTG |
| pMut1 sfGFP rev | AGGACTGAGCTAGCTGTCAAGTTTCAGTGGTGCGTACAATTAAG |
| pMut1 sacB gmR fwd | GCGCTGAACGCGATTCTG |
| pMut1 sacB gmR rev | GTTTCAGTGGTGCGTACAATTAAG |
| pMut2 sacB ampR fwd | GGCCTTATCTAAAACAGAC |
| pMut2 sacB ampR rev | CACTGCAACCCAACATTG |
| pEX18 sacB gmR fwd | CCTAAGCCCCAAAATATTTAAAGTATATATTATATGGTATATTCATACACATATACCTGCCGTTCAC |
| pEX18 sacB gmR rev | ATTGTACGCACCACTGAAACTTAGGTGGCGGTACTTGG |
| pEX100T sacB ampR fwd | GTCAGAATCGCGTTCAGCGCCACATATACCTGCCGTTC |
| pEX100T sacB ampR rev | GGCAGAAACATTAAATAATGACAATGTTGGGTTGCAGTGTTACCAATGCTTAATCAGTGAGGCACC |
| pKD4 kanR internal fwd | CGTTGGCTACCCGTGATATTG |
| pKD4 kanR internal rev | CGTCCTGCAGTTCATTCAGG |
| EcN clbQ ko fwd | CATCAAATTAAACGAATTCTATTACACAACAAGGAGTGGGACGATGAGTAAGTGTAGGCTGGAGCTGCTTC |
| EcN clbQ ko rev | GATGATGGAACAGCCATATCTATTGCTCCTTGTATAGTTACACAACTATTTCATATGAATATCCTCCTTAGTTCCTATTC |
| EcN clbQ internal fwd | CATTAACCTGTCCGATCG |
| EcN clbQ internal rev | GCCTGAAATACATACTGCTC |
| EcN clbQ external fwd | CACAACAAGGAGTGGGACGATG |
| EcN clbQ external rev | GGAACAGCCATATCTATTGCTCC |
| EcN fim ko fwd | GAACGACTGCCCATGTCGATTTAGAAATAGTTTTTTAAAGGAAAGCAGCGTAGGCTGGAGCTGCTTC |
| EcN fim ko rev | TAGCTTCAGGCAATATTGCGTACCTGCATTAGCAATGTTCTGTGATTTCTCATATGAATATCCTCCTTAGTTCCTATTC |
| EcN fim external fwd | CGTTTCTGTGGCTCGACG |
| EcN fim internal rev | CTAGCGGTACGAACCTGGC |
| EcN foc ko fwd | TGTGATGACAGATACGGTGTGCGTAGTTCAATTAAAAACAGGAATTAAATGTAGGCTGGAGCTGCTTC |
| EcN foc ko rev | GAAAAACTGGATATAAAGAGCAGTAATATCATTACCGCCACAACTGCATTCATATGAATATCCTCCTTAGTTCCTATTC |
| EcN foc external fwd | GCAGTCAAATGAGAGTGCGG |
| EcN foc internal rev | AGTGTTTACAGCACATGCAGC |
| EcN ecp ko fwd | AATGACTCACCGGGACTTCATGTCCTCAATTCAACTCGGGAAGAAAAGCAGTAGGCTGGAGCTGCTTC |

| | |
|---|---|
| EcN ecp ko rev | GATTTTTGAGTCTCTGAGGTGGAATTCTTCCCTCGTCCGATGGATAAGTCCATATGAATATCCTCCTTAGTTCCTATTC |
| EcN ecp external fwd | CGTGGTATACGCTGGACTGAG |
| EcN ecp internal rev | GCTACCGAGTGGCGTCAC |
| EcN csgBA ko fwd | GTTGAAATGATTTAATTTCTTAAATGTACGACCAGGTCCAGGGTGACAACGTAGGCTGGAGCTGCTTC |
| EcN csgBA ko rev | TTAAAGGTTATCTGACTGGAAAGTGCCGCAAGGAGTAATAACGCATTCATCATATGAATATCCTCCTTAGTTCCTATTC |
| EcN csgBA external fwd | CTTAAATGTACGACCAGGTCC |
| EcN csgBA internal rev | CGAATAGCCATTTGCGACTGTC |
| EcN flu ko fwd | CTGACCATGCTGTTTTGTACCTGCCGGTATCCACATTTGTGGGTACCGGCGTAGGCTGGAGCTGCTTC |
| EcN flu ko rev | GGGCAATAAACGTGATGTACAACCCGGCATATCTGTTGCTCCCTGAAAGTCATATGAATATCCTCCTTAGTTCCTATTC |
| EcN flu external fwd | CTGACCATGCTGTTTTGTAC |
| EcN flu internal rev | TTCAGAGTGGTATTCACTGCCTG |
| EcN bcsA ko fwd | GCCTGTTAAACTATTCCGGGCTGAAAACGCCAGTCGGGAGTGCATCATGAGTAGGCTGGAGCTGCTTC |
| EcN bcsA ko rev | CCAGAGCCACTGCACAAATCCAGAATATTTTTCTTTTCATCGCGTTATCACATATGAATATCCTCCTTAGTTCCTATTC |
| EcN bcsA external fwd | GGCTGAAAACGCCAGTCG |
| EcN bcsA internal rev | ACGATGATACCAAGGATCAACCG |
| EcN wcaLM ko fwd | GAATGCAGATGGTGCAATCTGTGCTTGAACGCATCGGGGAGGTGAAATGAGTAGGCTGGAGCTGCTTC |
| EcN wcaLM ko rev | GTAGCATTGTTCCTAAGTATGGCTCCATTTTTCCAGGAATGGTCGCAAATCATATGAATATCCTCCTTAGTTCCTATTC |
| EcN wcaLM external fwd | GAATGCAGATGGTGCAATC |
| EcN wcaLM internal rev | GCGAATGACACCCAGTTCG |
| EcN pga ko fwd | CTGTAATTAGATATAGAGAGAGATTTTGGCAATACATGGAGTAATACAGGGTAGGCTGGAGCTGCTTC |
| EcN pga ko rev | AGTGTATTATCGGCGCAGAGCCCGGGCGAACCGGGCTTTGTTTTGGGTGTCATATGAATATCCTCCTTAGTTCCTATTC |
| EcN pga external fwd | GGAATGGATTTTCGGGCGAG |
| EcN pga internal rev | ATTATTCCGTAATGCCTGCACG |
| EcN fliC ko fwd | GGTGGAAACCCAAAACGTAATCAACGACTTGCAATATAGGATAACGAATCGTAGGCTGGAGCTGCTTC |
| EcN fliC ko rev | ATCAGGCAATTTGGCGTTGCCGTCAGTCTCAGTTAATCAGGTTACGGCGACATATGAATATCCTCCTTAGTTCCTATTC |
| EcN fliC external fwd | GTGGAAACCCAAAACGTAATC |
| EcN fliC internal rev | CATTCAGCCCCAGCGTATC |

| | |
|---|---|
| EcN waaQ ko fwd | GCTTTTTCTGCTTCACCTTAATCGGATAATCTCAACAAAAAGAGTTACTTGTAGGCTGGAGCTGCTTC |
| EcN waaQ ko rev | TGCAAACCGCCAAAGGGAAAATATTTATATAAACAAAAAGCAACGATCATCATATGAATATCCTCCTTAGTTCCTATTC |
| EcN waaQ external rev | CTTTCTCCTGCGCTACTTTCT |
| EcN waaG ko fwd | CAGCTGTCGATAAATTACTTCCCTCCTCCACGACAGGTACGTCGTTATGAGTAGGCTGGAGCTGCTTC |
| EcN waaG ko rev | GGATCTTTACCGCGCCATAACGTGGCAAACGGCTCTTTAAGTTCAACCACATATGAATATCCTCCTTAGTTCCTATTC |
| EcN waaG external fwd | GCGTCCCTGGTCAAATAACA |
| EcN waaG internal rev | CTTTCTCCTGCGCTACTTTCT |
| EcN waaT ko fwd | TATTTATTATTTTTATTTAAAGATAATAAAATGACATTGAGAAAAGTTAAGTAGGCTGGAGCTGCTTC |
| EcN waaT ko rev | TAATAAAATGAGAAGCCGCGATAGCGTATACTTGTAATCATAAGAGTAATCATATGAATATCCTCCTTAGTTCCTATTC |
| EcN waaT external fwd | AAGCCGATATCCAGCGTTTAT |
| EcN waaT internal rev | CCGCCTGAGTACTATCATTGTC |
| EcN waaP ko fwd | GTCTGCCAGAGAAAGCGGCGGACATCATAACGGGTGGTCTGGATGGTTGAGTAGGCTGGAGCTGCTTC |
| EcN waaP ko rev | ATCATCTCTTGTGGATTAAAATAGTGGGCACTCATATTTCTCTCCGGAAACATATGAATATCCTCCTTAGTTCCTATTC |
| EcN waaP external fwd | CGTCGATAGAGTCGTTGGATTT |
| EcN waaP internal rev | CATCCCGCAGTCGATGAAT |
| EcN waaO ko fwd | AAATCAGGGAAAGAACGATTCGAAAATCGTTGTAATTTCCGGAGAGAAATGTAGGCTGGAGCTGCTTC |
| EcN waaO ko rev | AATACGAAAACCGTTCTTTTATAAATTCATTCATTTAACTTTTCTCAATGCATATGAATATCCTCCTTAGTTCCTATTC |
| EcN waaO external fwd | GGCATTAACCACCGTGACTG |
| EcN waaO internal rev | AGCAACATGGAATGATGATGCC |
| EcN waaY ko fwd | AAGGTATAACTTCATTAATTAAGTACAAGCTTAAGAAATAAATTACTCTTGTAGGCTGGAGCTGCTTC |
| EcN waaY ko rev | TAATACTCTCAGCTAATAAATCCATGTTGGTTCCGTTTTGACTGTGTGGTCATATGAATATCCTCCTTAGTTCCTATTC |
| EcN waaY external fwd | GATGCCGATGTCGTTTGTAAAG |
| EcN waaY internal rev | GGTCGCCCGATAGCATATTT |
| EcN waaW ko fwd | TAAAAAATTAAAAGGCAAAGCGTAAACCACACAGTCAAAACGGAACCAACGTAGGCTGGAGCTGCTTC |
| EcN waaW ko rev | ATAGTACTCATCCTTAATTATTATTGTAACTCAGACATCCATGATTTTTACATATGAATATCCTCCTTAGTTCCTATTC |
| EcN waaW external fwd | GGGCGTAGAGCTCAATGATATG |

| | |
|---|---|
| EcN waaW internal rev | CCACATCATTCCAGCAAGAAAC |
| EcN waaV ko fwd | AATTAAGGATGAGTACTATAATATTTTATTAGTTCAGGCAAGAGAACACGGTAGGCTGGAGCTGCTTC |
| EcN waaV ko rev | GACATATAATTTCAACCTATGCTACGAGGATGGGTTATTTAATGCAAGACCATATGAATATCCTCCTTAGTTCCTATTC |
| EcN waaV external fwd | CTGGAACGACTGGACGAATTAT |
| EcN waaV internal rev | CCGTCTCATGTTGTCTGGAATA |
| EcN waaL ko fwd | GTTAGGTCTTGCATTAAATAACCCATCCTCGTAGCATAGGTTGAAATTATGTAGGCTGGAGCTGCTTC |
| EcN waaL ko rev | TAATTTTGAAATAAAATCAGCTTCCTTGCTGATTTTATTTCACATATTCACATATGAATATCCTCCTTAGTTCCTATTC |
| EcN waaL external fwd | AAGGCGCAAGGACAGTTTA |
| EcN tatC ko fwd | GAACCGAAAACCGCTGCACCTTCCCCTTCGTCGAGTGATAAACCGTAAACGTAGGCTGGAGCTGCTTC |
| EcN tatC ko rev | ATCAAACATCCTGTACTCCATATGACAACCGCCCTGACGGGCGGTTGAATCATATGAATATCCTCCTTAGTTCCTATTC |
| EcN tatC external fwd | CTCCAGGAGTTTCAGGACAGTC |
| EcN tatC internal rev | GATCATCGTTGAACCTTGCGG |
| EcN manC ko fwd | TTGGGAATCAAAAGTTGCCAATTTAATGAATACATTAGATGAGAAATTATGTAGGCTGGAGCTGCTTC |
| EcN manC ko rev | ATTAACTAAAAATATTAGCGCTCTTCTACTAACGTTACATTATCAAAAATCATATGAATATCCTCCTTAGTTCCTATTC |
| EcN manC external fwd | GTTCGTCTTGGAGCAGGC |
| EcN manC internal rev | GGTTGGTACAATCCAAACCATCC |
| EcN mobA ko fwd | GAGACAGACACGTTAGCAGGGTCAATCCCACAATAAAAGAGGCGATATCGGTAGGCTGGAGCTGCTTC |
| EcN mobA ko rev | CCACTCCATGCAGCAAAGGCGAGTAACGGTATCATCGTTTTTCCTGCCATCATATGAATATCCTCCTTAGTTCCTATTC |
| EcN mobA external fwd | GGATCACCGCATCTTTCGC |
| EcN mobA internal rev | AGAAAGCTGCGTCATAAGCG |
| EcN motAB ko fwd | CGCCTGACGACTGAACATCCTGTCATGGTCAACAGTGGAAGGATGATGTCGTAGGCTGGAGCTGCTTC |
| EcN motAB ko rev | CTTCATCAAAAAATGTCTGATAAAAATCGCTTATATCCATGCTCACGCTGCATATGAATATCCTCCTTAGTTCCTATTC |
| EcN motAB external fwd | GGTGCGGTTTGTTGAAAGTGG |
| EcN motAB internal rev | GAAACAGCAACGGCAGCG |
| EcN ymgBC ko fwd | GAAAATAAACAGAATACATTAAAATTTCATAAGTAAGATGAGAGGTTACCGTAGGCTGGAGCTGCTTC |

| | |
|---|---|
| EcN ymgBC ko rev | CATCAGCATAGTGATACAGCTGATGTTTATTCAAAAACCT TACTCAAGTTCATATGAATATCCTCCTTAGTTCCTATTC |
| EcN ymgBC external fwd | GCGACAGGGGAAGAGGAAC |
| EcN ymgBC internal rev | ATTAAATTGGTCACAGCCTGCC |
| EcN 10090-10165 ko fwd | CCACTGCTCTCGCGAATAAAGATGGAAAATCAATCTCATG GTAATAGTCCGTAGGCTGGAGCTGCTTC |
| EcN 10090-10165 ko rev | TCAGTGCCCTGCGCTGTTTTTCGCGTGCGCCAGGTAAACG TGTCCGTCACCATATGAATATCCTCCTTAGTTCCTATTC |
| EcN 10090-10165 external fwd | CAGAGTATCCTTCAGGCTCTGG |
| EcN 10090-10165 internal rev | CCCATCTATCCCATATCCAGCG |
| EcN 10175-10190 ko fwd | GATCCCCTTCAATCACACACAGCGCCATCCGAACTATCGG AGGTGAGGCTGTAGGCTGGAGCTGCTTC |
| EcN 10175-10190 ko rev | CGGGCTGTTGCATTATCACAGGCACTCAGTGAATGCCTGC TGTAATGCCGCATATGAATATCCTCCTTAGTTCCTATTC |
| EcN 10175-10190 external fwd | CATCCATGCGAAAAACCTGCC |
| EcN 10175-10190 internal rev | CAGTCGCTCAGGCTTAACCC |
| EcN yhhT qPCR fwd | GCCATTACGATTGTGGTTGTG |
| EcN yhhT qPCR rev | CCTCGGCAGCATAGAGATAAAT |
| EcN ymgC qPCR fwd | AGATGAGACCAGTGTTATTCTTTCT |
| EcN ymgC qPCR rev | GAGCACAGATTCCCTGTCATTA |
| EcN 10090 qPCR fwd | CACTGTGTCCTCCTCGAATAAA |
| EcN 10090 qPCR rev | GGGAGTGACGGAAGCATAAA |
| EcN 10145 qPCR fwd | GCATCGGTCTGGAGAAGATAAA |
| EcN 10145 qPCR rev | TCAGGCGCAGCAAACTTA |
| EcN 10190 qPCR fwd | CAACATCCAGGACACCTCTTT |
| EcN 10190 qPCR rev | GCACCATTTACAGCAGACTTTC |
| EcN lrhA qPCR fwd | CATTTCTGCCATGTAGGCATTAC |
| EcN lrhA qPCR rev | TGCCGATACGATCTTACCTTTC |
| EcN flgB qPCR fwd | GCCTTCGCTTGACGGTAATA |
| EcN flgB qPCR rev | CGCTAAGGCTCATCTGGTATTG |
| EcN frr qPCR fwd | GCAAGCGTAACGGTAGAAGA |
| EcN frr qPCR rev | CCAAGATCGGACGCCATAAT |
